# Evolution of increased longevity and slowed ageing in a genus of tropical butterfly

**DOI:** 10.1101/2025.08.29.673072

**Authors:** Jessica Foley, Josie McPherson, Made Roger, Cruz Batista, Rémi Mauxion, Greta Hernández, Richard Kelson, Fletcher J. Young, W. Owen McMillan, Stephen H. Montgomery

## Abstract

Evolution has given rise to lifespans in extant species ranging from days to centuries. Given that mechanisms of ageing are highly conserved, studying long-lived lineages across the animal kingdom could yield insights relevant for healthy ageing in humans. However, typical models of extended lifespan often live for decades, making them impractical for longitudinal studies. Ideal model systems would be organisms that are naturally long-lived compared to their close relatives, but have lifespans on experimentally tractable scales. Here, we present the Neotropical butterfly genus *Heliconius* as a novel model system for the evolution of extended longevity. We collate data from 27 species across the Heliconiini tribe to reveal a 25-fold variation in lifespan within the group, with our 348-day maximum for *Heliconius hewitsoni* longer than any butterfly species previously recorded in the scientific literature. While previous work has attributed this lifespan extension to a plastic response to enhanced nutrition, we conduct detailed survival and functional senescence analyses on two species representative of shorter- and longer-lived clades to show evidence of evolved, heritable mechanisms of slowed ageing in *Heliconius*. Our results add a new case study to the canon of noteworthy agers, and provide valuable insights into the evolution of increased longevity.

## Introduction

The diversity of lifespans across the animal kingdom, ranging from days to centuries [1], has intrigued scholars for millennia [2]. More recent work on age-specific mortality has now shown that variation in rates of ageing is just as pronounced; while humans show steep increases in mortality with age, for many species these inclines are much gentler [3].

Evolutionary explanations for senescence, defined as the physiological deterioration that accompanies ageing, rest on the concept of the “selection shadow”, the declining force of natural selection with age [4–7]. This effect is thought to provide an explanation for the prevalence of declines in physiological function and survival with age. Yet the observed diversity in ageing across the tree of life [3] suggests that many organisms have evolved adaptations allowing them to slow or delay senescence. Considering the high evolutionary conservation of mechanisms of ageing [8], uncovering such adaptations across diverse taxa may provide unique insights for the study of healthy ageing.

As the most species-rich animal class, insects are famed for their morphological and ecological diversity. They also display extreme variation in longevity, with maximum lifespans ranging from just a few days in adult mayflies to up to several decades in the case of some ant and termite reproductive castes. This represents a 5000-fold difference within the class – versus, for comparison, the 60-fold difference in lifespan observed in mammals [9]. The majority of research on insect ageing has thus far focused on intra-species differences, in particular in the fruit fly, *Drosophila melanogaster*, where the vast genetic toolbox has allowed for the elucidation of specific genes regulating lifespan [10, 11]. Similarly, research on the dramatic differences in lifespan between social insect castes has provided valuable insights into the molecular underpinnings of such plasticity [12, 13]. This work has been instrumental in uncovering the proximate mechanisms of lifespan extension. However, leveraging the rich inter-specific diversity in insect lifespan, in combination with the depth of available ecological knowledge, can help to illuminate the evolutionary origins of lengthened life. In particular, inter-species lifespan differences within insects can be long in relative terms but short in absolute terms, making them much more tractable for longitudinal studies than typical mammalian and avian models of extended longevity.

The long lifespans of the genus *Heliconius* rank among the longest recorded in butterflies [14], and have been reported to stretch to at least 6 months in the wild [15]. This represents a dramatic lifespan extension over the ∼6 weeks reported for their close relatives in the Heliconiini tribe [16], from whom they diverged relatively recently in evolutionary time (∼18mya) [17]. The remarkable increase in *Heliconius’* longevity has been linked to the evolution of adult pollen-feeding in this genus, which is unique among butterflies [16, 18, 19]. Pollen-feeding provides *Heliconius* with a source of amino acids in adulthood, in contrast with their Heliconiini relatives, which as adults must rely on nitrogenous resources derived from larval phytophagy [18, 20]. All *Heliconius* species actively collect and ingest pollen, with the exception of one clade of 4 species, whose phylogenetic position has been debated [17, 18]. This dietary innovation has been linked to a suite of other behavioural, neuroanatomical, and physiological traits in *Heliconius* that are not present in other non-pollen-feeding Heliconiini [18]. These include increased investment in neural structures supporting learning and memory [21] and more stable long-term visual memories [22]. Importantly, pollen-derived amino acids have been shown to facilitate a lengthened reproductive lifespan in female *Heliconius*, allowing for the continual production of oocytes throughout adulthood [16, 23], whereas pollen-deprived *Heliconius* and their non-pollen-feeding Heliconiini relatives instead exhibit rapid reproductive senescence beginning at ∼3 weeks of age [16]. Pollen-consumption has also been suggested to directly extend *Heliconius’* lifespan, with pollen-deprived *Heliconius* reportedly living no longer than their non-pollen-feeding relatives [16]. However, the evidence for a direct effect on longevity is more equivocal, and later pollen-deprivation experiments have not recapitulated these results [23, 24]. The suite of other derived traits linked to pollen-feeding in *Heliconius* [18] instead suggests that across evolutionary time, enhanced nutrition may have indirectly facilitated longevity in this genus via an investment in longer-lived tissues. The lengthened reproductive lifespan facilitated by this nutritional advantage would likely favour selection for longevity due to the enhanced fitness of long-lived, continually reproducing individuals.

Despite *Heliconius’* potential as a model for the evolution of longevity, available data is restricted to scattered observations of maximum lifespan. These data point to significant increases in *Heliconius* over their non-pollen-feeding Heliconiini relatives (see Table 1). However, discussion of this lifespan extension tends to refer to a single broad shift in *Heliconius*, despite observed variation in the degree of this increase between *Heliconius* species. Maximum lifespan is also a notoriously problematic metric of ageing, correlating highly with sample size and being unrepresentative of an average individual [25, 26]. More thorough survival data from a wider range of species across the tribe would allow for estimates of more reliable metrics such as median lifespan, which are less sensitive to sample size [27]. Parametric survival analysis, which fits survival data to a hazard function to understand mortality patterns over time, can also help to disentangle the basis of observed lifespan differences by offering insights into how not just survival, but also rates of ageing, vary between populations [25]. These techniques allow for the identification and quantification of actuarial senescence, defined as an increase in mortality with age. This is distinct from physiological senescence, which measures the decline in physiological function that may often accompany the ageing process, although actuarial senescence may imply the presence of physiological senescence [28]. A characterisation of physiological senescence in this system would also allow for a thorough interrogation of the direct and indirect impacts of pollen-feeding on *Heliconius* life history, and provide a reliable index upon which future ageing-related interventions may be based. To date, the only description of physiological senescence in the Heliconiini tribe is limited to the reproductive senescence demonstrated by *Dryas iulia* and pollen-deprived *Heliconius charithonia* [16].

**Table 1:** Maximum reported lifespans for species in the Heliconiini tribe.^1^.

| Species | Feeding habit | Maximum lifespan (days) | Data source |
| --- | --- | --- | --- |
| *Heliconius hewitsoni* | pollen-feeding | 348 | Butterfly house (unpublished) |
| *Heliconius hecale* | pollen-feeding | 277 | Butterfly house (unpublished) |
| *Heliconius erato* | pollen-feeding | 274 | Butterfly house (unpublished) |
| *Heliconius melpomene* | pollen-feeding | 260 | Butterfly house (unpublished) |
| *Heliconius ismenius* | pollen-feeding | 245 | Butterfly house (unpublished) |
| *Heliconius cydno* | pollen-feeding | 244 | Butterfly house (unpublished) |
| *Heliconius atthis* | pollen-feeding | 210 | Butterfly house (unpublished) |
| *Heliconius numata* | pollen-feeding | 210 | Butterfly house (unpublished) |
| *Heliconius hortense* | pollen-feeding | 198 | Butterfly house (unpublished) |
| *Heliconius charithonia* | pollen-feeding | 184 | Butterfly house (unpublished) |
| *Heliconius sapho* | pollen-feeding | 177 | Butterfly house (unpublished) |
| *Heliconius sara* | pollen-feeding | 170 | Butterfly house (unpublished) |
| *Heliconius ethilla* | pollen-feeding | 162 | Mark-release-recapture [15] |
| *Heliconius clysonymus* | pollen-feeding | 156 | Butterfly house (unpublished) |
| *Heliconius hermathena* | pollen-feeding | 139 | Mark-release-recapture [64] |
| *Heliconius pachinus* | pollen-feeding | 106 | Butterfly house [65] |
| *Dryas iulia* | non-pollen-feeding | 98 | Butterfly house (unpublished) |
| *Heliconius hecalesia* | pollen-feeding | 89 | Butterfly house (unpublished) |
| *Heliconius eleuchia* | pollen-feeding | 86 | Butterfly house (unpublished) |
| *Dryadula phaetusa* | non-pollen-feeding | 85 | Butterfly house (unpublished) |
| *Heliconius nattereri* | pollen-feeding | 77 | Mark-release-recapture [66] |
| *Heliconius doris* | pollen-feeding | 69 | Butterfly house (unpublished) |
| *Heliconius xanthocles* | pollen-feeding | 66 | Mark-release-recapture [67] |
| *Agraulis vanillae* | non-pollen-feeding | 60 | Butterfly house [65] |
| *Eueides isabella* | non-pollen-feeding | 46 | Butterfly house (unpublished) |
| *Philaethria dido* | non-pollen-feeding | 44 | Butterfly house [65] |
| *Dione juno* | non-pollen-feeding | 14 | Insectary population [68] |

<sup>1</sup> “Mark-release-recapture” refers to studies conducted on wild populations; “insectary population” refers to studies conducted on captive populations reared in insectary conditions (typically in smaller cages); “butterfly house” refers to studies conducted on captive populations living in semi-natural conditions in commercial butterfly houses. The single highest record for each species is shown; for additional records, see Supplementary Data 1.

To clarify *Heliconius’* utility as a model system for the evolutionary and mechanistic bases of ageing, it is important to determine whether longevity in this genus reflects a simple plastic nutritional advantage, or is instead indicative of evolved, genetically determined traits which may offer more wide ranging insights into longevity. Here, we leverage the neat experimental framework provided by comparisons with the closely-related but shorter-lived Heliconiini outgroups. We assess the impact of pollen consumption on lifespan and senescence in butterflies in this tribe, to better understand the benefits of pollen-feeding to evolutionary fitness. We collate data from multiple complementary sources to build a profile of ageing across the Heliconiini tribe, broadly supporting a shift towards longer lifespans co-occurring with the transition to a pollen-feeding behaviour, but revealing diversity in the parameters of ageing within the *Heliconius* genus. We then narrow our focus to one representative long-lived pollen-feeder, *Heliconius hecale*, and one representative short-lived non-pollen-feeder, *Dryas iulia*, combining survival analyses and indices of physiological senescence under both pollen-fed and pollen-deprived diet treatments to disentangle the impact of diet on *Heliconius* ageing and lifespan. Our results provide the first evidence for an evolved lifespan extension and slowed ageing in this genus, establishing *Heliconius* as a valuable new case study for the evolution of long life.

## Methods

More detailed methodology may be found in the Supplementary Information.

### Heliconiini maximum reported lifespan data collation

Data on maximum reported lifespan for species across the Heliconiini tribe (Table 1, Supplementary Data 1) was collated from a literature search within PubMed, Google Scholar, and Web of Science for long-term studies of *Heliconius* and other Heliconiini butterflies, using combinations of the search terms “*Heliconius*”, “Heliconiini”, “lifespan”, “longevity”, “mark”, “recapture”, and “population dynamics”, between January and April 2021. However, considering the popularity of Heliconiini species in commercial butterfly houses, active exhibits listed on the International Association of Butterfly Exhibitors and Suppliers were also contacted to enquire if they had collected lifespan data, as has been done previously [29]. For many species, multiple records of maximum longevity were found, sourced from data from butterfly exhibitors, mark-release-recapture studies, and insectary populations (Supplementary Data 1). In all such instances, maximum longevity records from butterfly exhibitors eclipsed those from insectary populations and mark-release recapture studies in the wild.

### Butterfly husbandry and survival data collection

All butterflies were captive-reared from stock populations at the Smithsonian Tropical Research Institute (STRI) outdoor insectaries in Gamboa, Panama. All Panamanian butterflies were collected under permit number SE/A-14-18 and SE/A-82-19.

#### Survival data from a multi-species cognitive experiment cohort

Survival data was collected for *Heliconius hecale melicerta, Heliconius melpomene rosina*, *Dryadula phaetusa*, and *Agraulis vanillae* during a previous cognitive experiment published in [22]. Survival data for *Dryas iulia* was obtained during a similar cognitive experiment (Foley *et al*., in prep) conducted under the same conditions. Newly-eclosed butterflies were placed into experimental cages and fed with a sucrose-protein solution (25% w/v sucrose and 5% w/v Vetark Critical Care Formula), replaced daily. Butterflies were subjected to long-term memory assays, further details of which may be found in [22], and maintained until the end of their natural lifespans. In brief, during these assays butterflies were kept in free flight cages (2m [L] x 3m [W] x 2m [H]), and were fed *ad libitum*, except prior to or during a series of preference assays (see Supplementary Methods for further details). Cages were checked daily for dead individuals, death dates were recorded, and missing or predated individuals were censored at the age at which they were last seen alive. In total, survival data from both sexes was analysed for 175 individuals of *A. vanillae*, 263 individuals of *D. iulia*, 108 individuals of *D. phaetusa*, 120 individuals of *H. hecale*, and 103 individuals of *H. melpomene*.

#### Semi-natural “mark-release-recapture” cohort

Newly-eclosed butterflies of 20 different species were marked with a unique ID and released into a large 11m (L) x 11m (W) x 6m (H) cage filled with trees, *Passiflora* host-plants, and other flowering plants, creating a semi-natural environment. Data collection was carried out on a weekly basis, with some omissions due to time constraints. Approximately 15 minutes were spent patrolling the cage and recording the IDs of any butterflies visually identified and alive on that date. To ensure minimal intervention, butterflies were not recaptured or handled. In total, 959 butterflies of 20 different Heliconiini species were released into the cage, including: *A. vanillae* (n = 12), *D. iulia* (n = 34), *Dione juno* (n = 45), *D. phaetusa* (n = 30), *Eueides isabella* (n = 59), *Heliconius atthis* (n = 33), *Heliconius charithonia* (n = 3), *Heliconius cydno* (n = 19), *Heliconius doris* (n = 32), *Heliconius erato* (n = 46), *Heliconius hewitsoni* (n = 15), *H. hecale* (n = 47), *Heliconius himera* (n = 2), *Heliconius ismenius* (n = 20), *H. melpomene* (n = 285), *Heliconius numata* (n = 94), *Heliconius pachinus* (n = 19), *Heliconius sapho* (n = 81), *Heliconius sara* (n = 70), and *Philaethria dido* (n = 13). Species selected were those being reared at the STRI insectaries at the time of the experiment. All *Heliconius* included in this study are pollen-feeding species, and data for the four non-pollen feeding *Heliconius* in the Aoede clade are unfortunately not available. This reflects the understudied nature of these species, which occur at low densities and have a derived host plant, and are therefore challenging to work with [30]. Sample number reflects availability of pupae. Both sexes were represented for all species.

#### Pollen-manipulation experiment cohort

Newly-eclosed individuals of *H. hecale* and *D. iulia*, selected for their local abundance and ease of rearing, were sexed and weighed, their forewings were measured, and they were marked with a unique ID. They were then randomly assigned to either a pollen-fed or pollen-deprived treatment. Pollen-fed cages contained flowering plants to serve as natural pollen sources as well as artificial feeders filled with a 20% w/v sucrose and 10% w/v organic, pesticide-free bee pollen (Bienenschwarmmm & Aspermühle) solution. Pollen-deprived cages contained non-flowering plants as well as artificial feeders filled with a 20% w/v sucrose solution. The number of artificial feeders in this cage was increased to match the number of flowers + feeders in the pollen-fed cage to ensure feeding opportunities were as standardised as possible between cages. Cages were checked daily for dead individuals, death dates were recorded, and missing or predated individuals were censored at the age at which they were last seen alive. In total, survival data from both sexes was collected across the lifespan of 96 individuals of *H. hecale* (*n* _pollen-fed_ = 47, *n* _pollen-deprived_ = 49) and 116 individuals of *D. iulia* (*n* _pollen-fed_ = 57, *n* _pollen-deprived_ = 57).

### Functional senescence assays

Butterflies from the pollen-manipulation experiment cohort (*n _H. hecale_* = 96; *n _D. iulia_* = 116) were measured every two weeks for indices of functional senescence. Considering evidence from several insects for age-related declines in body mass [31, 32], and muscle function [33], butterflies were first weighed using a Sartorius Entris balance and then assayed for grip strength using a method adapted from [34]. Grip strength provides a proxy for whole organism condition in butterflies [34], beetles [35], and is a standard biomarker of health in humans [36]. A custom-built device consisting of a perch mounted on a lightweight base was placed on the balance, which was then tared. Butterflies were held by their wings and allowed to grasp the perch, then gently pulled upwards until they released it. The peak negative reading on the balance from this exercise was taken as a measure of grip strength, which we took to be a proxy for muscle function. This was repeated five times for each individual, and the maximum reading was used for statistical analysis as an indicator of the individual’s maximum capacity. Data on flight behaviour was also recorded and is presented in Supplementary Note 1.

### Statistical analyses

All statistical analyses were conducted using R v4.3.1 [37]. Non-parametric and semi-parametric survival analyses for the pollen-manipulation experiment and multi-species cognitive experiment cohorts were conducted with the aid of the packages *survival* v3.5-5 [38] and *coxme* v2.2-18 [39]. Cox proportional hazards models were created for each species including adult eclosion mass, diet, and sex (pollen-manipulation experiment cohort) or just sex (multi-species cognitive experiment cohort) as fixed effects. The package *flexsurv* v2.2.2 [40] was then used to create parametric survival models for these cohorts fit to a Gompertz distribution (see Supplementary Note 2 for an explanation of distribution selection). The Gompertz function models how mortality risk changes with age, and is described by the equation 𝜇(𝓍) = 𝛼 𝑒^𝛽𝓍^, where: 𝜇(𝓍) is the instantaneous mortality rate at age 𝓍, 𝛼 is the baseline mortality independent of age, and 𝛽 is the age-dependent mortality rate (i.e. the relative change in mortality with age 𝓍), also known as the rate of ageing [25, 41]. A value of 𝛽 > 0 reflects an increase in mortality risk with age, confirming the presence of actuarial senescence, and reflecting the process of ageing. Graphically speaking, when age is plotted against the natural log (ln) of the mortality rate, the intercept is equal to ln 𝛼 and the slope is equal to 𝛽, facilitating easy interpretation of these parameters: a higher intercept reflects an increase in baseline mortality risk, and a higher slope reflects an increased rate of ageing. Models allowed species (multi-species cognitive experiment cohort) or species and diet (pollen-manipulation cohort) to predict 𝛼 – baseline mortality, 𝛽 – rate of ageing, or both. The model with the lowest AIC was used for interpretation. Bootstrapped estimates (1,000 iterations) were generated for 𝛼 and 𝛽 for each group of interest, and the mean and 95% confidence intervals were extracted from these distributions. Estimates for these parameters were considered to be significantly different between groups if their 95% confidence intervals did not overlap.

Longevity data from the semi-natural “mark-release-recapture” cohort was analysed using the R package *BaSTA* v1.9.5 [42], which facilitates the use of incomplete recapture data to provide estimates of age-specific survival under a Bayesian framework. Models for each species were fit to a simple Gompertz distribution, running four parallel *BaSTA* simulations with 11,000 iterations, a burn-in of 1001, and a thinning rate of 200, to minimise serial auto-correlation. These models were then used to derive estimates for median lifespan as well as 𝛼 and 𝛽 parameters. Correlation between median lifespan estimates from *BaSTA* and existing reported maximum lifespans (Table 1) was performed using a Pearson’s correlation test. Results from this cohort and the multi-species cognitive experiment cohort, as well as existing reported maximum lifespans (Table 1), were then used to assess broad differences in ageing parameters between pollen-feeders (*Heliconius* species) versus non-pollen-feeders (the outgroup Heliconiini). Differences in median and maximum lifespans were assessed using two-tailed *t-*tests and differences in 𝛼 and 𝛽 Gompertz parameters for the semi-natural “mark-release-recapture” cohort were assessed using two-tailed Mann-Whitney U tests. To control for phylogenetic relatedness, these comparisons were then repeated using the phylANOVA() function from the R package *phytools* v2.3-0 [43], run with 10,000 simulations using a trimmed phylogenetic tree taken from [17]. This package and tree were also used to create the phylogenetic tree in Fig. 1, and to estimate phylogenetic signal as measured by Pagel’s λ. However, as this evolutionary transition only occurred once, we expect our results to be confounded by phylogeny, with limited power to disentangle these effects. As such, results from these phylogenetic ANOVAs are presented in Supplementary Note 3.

**Fig. 1:**
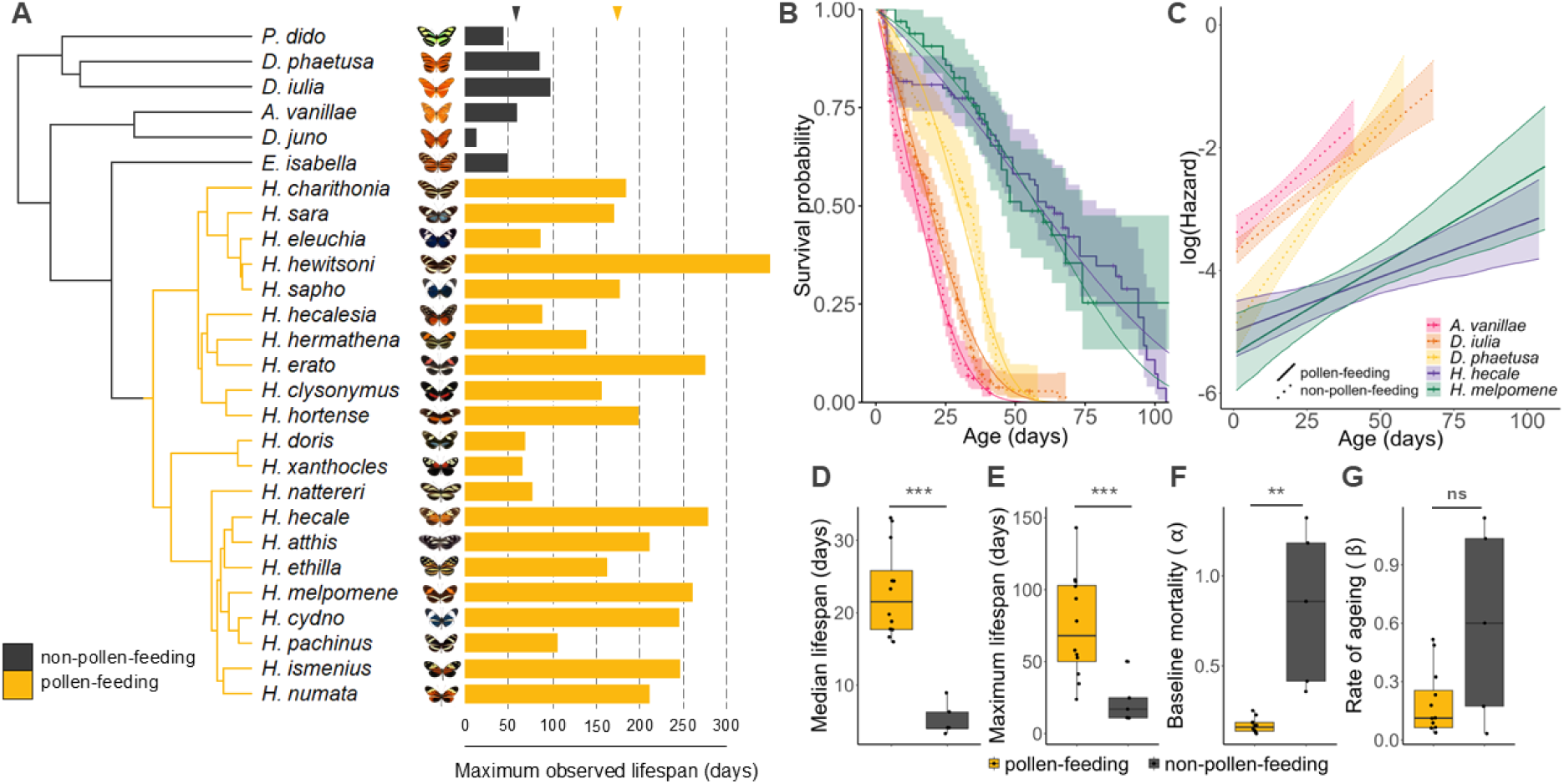
Pollen-feeding *Heliconius* spp. have longer lifespans and reduced ageing parameters relative to non-pollen feeding relatives. **A** Phylogeny of the Heliconiini tribe with associated bars representing maximum reported lifespans based on Tables 1 and 3. Branches are coloured according to feeding habit. Arrowheads indicate mean maximum lifespan for each feeding habit. **B** Kaplan-Meier survival estimates and 95% confidence intervals overlaid with the corresponding parametric survival curves for the multi-species cognitive experiment cohort. “+” indicates a censored data point. **C** log(Hazard) (equivalent to log[likelihood of death]) curves and 95% confidence intervals resulting from the transformation of the parametric survival function shown in **B** to the hazard function. An increase in the intercept on this graph represents an increase in baseline mortality (α); an increase in the slope represents an increase in the rate of ageing (β). Boxplots represent overall differences between pollen-feeders and non-pollen-feeders from the semi-natural “mark-release-recapture” cohort in **D** median lifespan estimates, **E** maximum observed lifespan, F baseline mortality estimates, and **G** rate of ageing estimates. Indications of statistical significance for these boxplots are taken from results of standard statistical techniques, without controlling for phylogenetic relatedness; see Supplementary Note 3 for results of phylogenetic ANOVA. n.s.: *p* > 0.05; *: *p* < 0.05; **: *p* < 0.01; ***: *p* < 0.001.

Body mass and grip strength data were analysed with LMMs using the package *lme4* v1.1-34 [44], for both single-species and interspecific datasets. All models included diet, age, and species (for the interspecific models) as fixed effects, as well as longevity to account for selective disappearance. All models also included individual ID as a random effect. Details of additional candidate predictors may be found in the Supplementary Information. An ANOVA was then run on the final model to report significance of named predictors.

## Results

### Reduced ageing parameters in pollen-feeding *Heliconius* species

Data collated from published field studies and public greenhouses reveals a 25-fold range in maximum reported lifespan for species across the Heliconiini tribe, from 14 days in *Dione juno* to an observed maximum of 348 days in *Heliconius hewitsoni* (Fig. 1, Table 1). Pollen-feeding *Heliconius* species have longer maximum reported lifespans, with a mean of 178.42 days as compared to 57.67 days for non-pollen-feeding Heliconiini outgroups (*t*_25_ = 5.69, *p* < 0.001). The phylogenetic signal for median lifespan is high (Pagel’s λ = 0.72), such that phylogenetic regression confounds the statistical significance of the group difference (Supplementary Note 3), consistent with a major effect of the transition to pollen-feeding in *Heliconius*.

Survival data from a multi-species cohort exposed to a cognitive experiment [22] also revealed higher lifespans in pollen-feeding *Heliconius* species (Fig. 1, Table 2), with a mean maximum lifespan of 105 days for pollen-feeders as compared with 55.67 days for non-pollen-feeders (*t*_3_ = – 4.84, *p* = 0.017), and a mean median lifespan of 56.5 days for pollen-feeders as compared with 24 days for non-pollen-feeders (*t*_3_ = – 4.38, *p* = 0.022). Sex was not a significant predictor of survival in any species (*D. phaetusa*: χ^2^ = 0.54, *p* = 0.464; *D. iulia*: χ^2^ = 1.43, *p* = 0.231; *H. hecale*: χ^2^ = 0.24, *p* = 0.627; *H. melpomene*: χ^2^ = 0.02, *p* = 0.884) with the exception of *A. vanillae* (χ^2^ = 13.38, *p* < 0.001). However, this exception appeared to be due to disproportionately high early mortality in *A. vanillae* males, as sex was no longer a significant predictor of survival when the population was subset to those that survived at least up to 5 days (χ^2^ = 2.87, *p* = 0.090). Beyond lifespan, parametric survival analysis of the multi-species cognitive experiment cohort showed differences between species in both baseline mortality (𝛼) and the rate of ageing (𝛽), with the best-supported model allowing both of these parameters to vary by species. Both of these parameters suggested reduced ageing in pollen-feeding *Heliconius* (Fig. 1, Table 2), though these did not always exceed formal statistical thresholds of significance (see Supplementary Note 4).

**Table 2:** Ageing parameters for the multi-species cognitive experiment cohort.

| Species | Feeding habit | Median lifespan (days) | Maximum lifespan (days) | Baseline mortality ($\alpha$) | Rate of ageing ($\beta$) |
| --- | --- | --- | --- | --- | --- |
| *Agraulis vanillae* | non-pollen-feeding | 17 | 41 | 0.033 | 0.044 |
| *Dryadula phaetusa* | non-pollen-feeding | 34 | 58 | 0.0075 | 0.067 |
| *Dryas iulia* | non-pollen-feeding | 21 | 68 | 0.024 | 0.040 |
| *Heliconius hecale* | pollen-feeding | 61 | 104 | 0.0070 | 0.018 |
| *Heliconius melpomene* | pollen-feeding | 52 | 106 | 0.0050 | 0.029 |

Data from a much larger sample of species across the Heliconiini tribe provided further evidence of reduced ageing in *Heliconius*. Results from Bayesian survival trajectory analysis of the semi-natural “mark-release-recapture” cohort showed pollen-feeding *Heliconius* to have higher maximum (*t*_15_ = 3.05, *p* = 0.008) and median lifespans (*t*_15_ = 6.02, *p* < 0.001) as well as reduced baseline mortality (𝛼) (*W* = 0, *p* < 0.001) compared with non-pollen-feeding outgroups (Fig. 1, Table 3, Supplementary Note 5). Rate of ageing (𝛽) is generally slower in pollen-feeding *Heliconius*, but this difference is not significant (*W* = 17, *p* = 0.195) (Fig. 1, Table 3, Supplementary Note 5). Median lifespan in this cohort appears to be significantly underestimated compared to the more thorough survival data from the multi-species cognitive experiment cohort. However, median lifespan still showed a significant positive correlation with maximum reported lifespans from Table 1 (Supplementary Note 5; *r* = 0.64; *t*_15_ = 3.25, *p* = 0.005).

**Table 3:** Ageing parameters for the semi-natural “mark-release-recapture” cohort.^2^.

| Species | Feeding habit | Median lifespan (days) | Maximum lifespan (days) | Baseline mortality ($\alpha$) | Rate of ageing ($\beta$) |
| --- | --- | --- | --- | --- | --- |
| *Agraulis vanillae* | non-pollen-feeding | 6.27 | 11 | 0.42 | 0.16 |
| *Dione juno* | non-pollen-feeding | 4.03 | 11 | 0.86 | 0.15 |
| *Dryadula phaetusa* | non-pollen-feeding | 8.93 | 17 | 0.36 | 0.086 |
| *Dryas iulia* | non-pollen-feeding | 3.33 | 25 | 1.32 | 0.025 |
| *Eueides isabella* | non-pollen-feeding | 4.03 | 50 | 1.18 | 0.0047 |
| *Heliconius atthis* | pollen-feeding | 16.63 | 53 | 0.25 | 0.016 |
| *Heliconius charithonia* | pollen-feeding | - | 22 | - | - |
| *Heliconius cydno* | pollen-feeding | 19.78 | 55 | 0.19 | 0.026 |
| *Heliconius doris* | pollen-feeding | 17.68 | 35 | 0.14 | 0.07 |
| *Heliconius erato* | pollen-feeding | 33.08 | 94 | 0.12 | 0.0092 |
| *Heliconius hecale* | pollen-feeding | 18.73 | 78 | 0.23 | 0.013 |
| *Heliconius hewitsoni* | pollen-feeding | 17.68 | 41 | 0.17 | 0.046 |
| *Heliconius himera* | pollen-feeding | -- | 17 | - | - |
| *Heliconius ismenius* | pollen-feeding | 15.93 | 24 | 0.15 | 0.074 |
| *Heliconius melpomene* | pollen-feeding | 30.28 | 102 | 0.14 | 0.0087 |
| *Heliconius numata* | pollen-feeding | 23.28 | 106 | 0.17 | 0.016 |
| *Heliconius pachinus* | pollen-feeding | 24.33 | 58 | 0.13 | 0.033 |
| *Heliconius sapho* | pollen-feeding | 32.73 | 143 | 0.13 | 0.0056 |
| *Heliconius sara* | pollen-feeding | 24.33 | 107 | 0.19 | 0.0053 |
| *Philaethria dido* | non-pollen-feeding | - | 7 | - | - |

<sup>2</sup> *H. charithonia*, *H. himera*, and *P. dido* were excluded from Bayesian survival trajectory analysis due to low sample size and so only maximum lifespan was collected for these species.

### Pollen consumption does not account for the full lifespan extension in *Heliconius*

We next conducted detailed survival analyses in two focal Heliconiini species to test the effects of pollen on longevity: *H. hecale*, as a representative pollen-feeding *Heliconius*, and *D. iulia,* as a representative non-pollen-feeding outgroup. Diet (i.e., provision of pollen) was a significant predictor of survival in *H. hecale* (χ^2^ = 8.86, *p* = 0.003), with a median survival of 47 days (maximum: 106 days) for the pollen-deprived group and a median survival of 63 days (maximum: 119 days) for the pollen-fed group (Fig. 2, Table 4). In contrast, diet was not a significant predictor of survival in *D. iulia* (χ^2^ = 0.34, *p* = 0.562), with a median survival of 29 days (maximum: 50 days) for the pollen-deprived group and a median survival of 27 days (maximum: 48 days) for the pollen-fed group (Fig. 2, Table 4). This is lower than both diet groups in *H. hecale*, implying both diet-dependent and diet-independent increases in longevity in *Heliconius*. Neither sex (*H. hecale*: χ^2^ = 0.73, *p* = 0.393; *D. iulia*: χ^2^ = 1.58, *p* = 0.208) nor eclosion mass (*H. hecale*: χ^2^ = 0.73, *p* = 0.530; *D. iulia*: χ^2^ = 0.16, *p* = 0.693) were found to be significant predictors of survival in either species. Parametric survival analysis showed that this reduced lifespan in pollen-deprived *H. hecale* was due to an increase in baseline mortality (𝛼), which was 1.98 times higher than that of the pollen-fed group, while the rate of ageing (𝛽) remained unchanged in pollen-deprived individuals, with the the best-supported model allowing 𝛼, but not 𝛽, to vary by diet (Fig. 2, Table 4). In accordance with its lack of effect on *D. iulia* lifespan, diet also had no effect on either baseline mortality (𝛼) or rate of ageing (𝛽) in this species (Fig. 2, Table 4; see Supplementary Note 6).

**Fig. 2:**
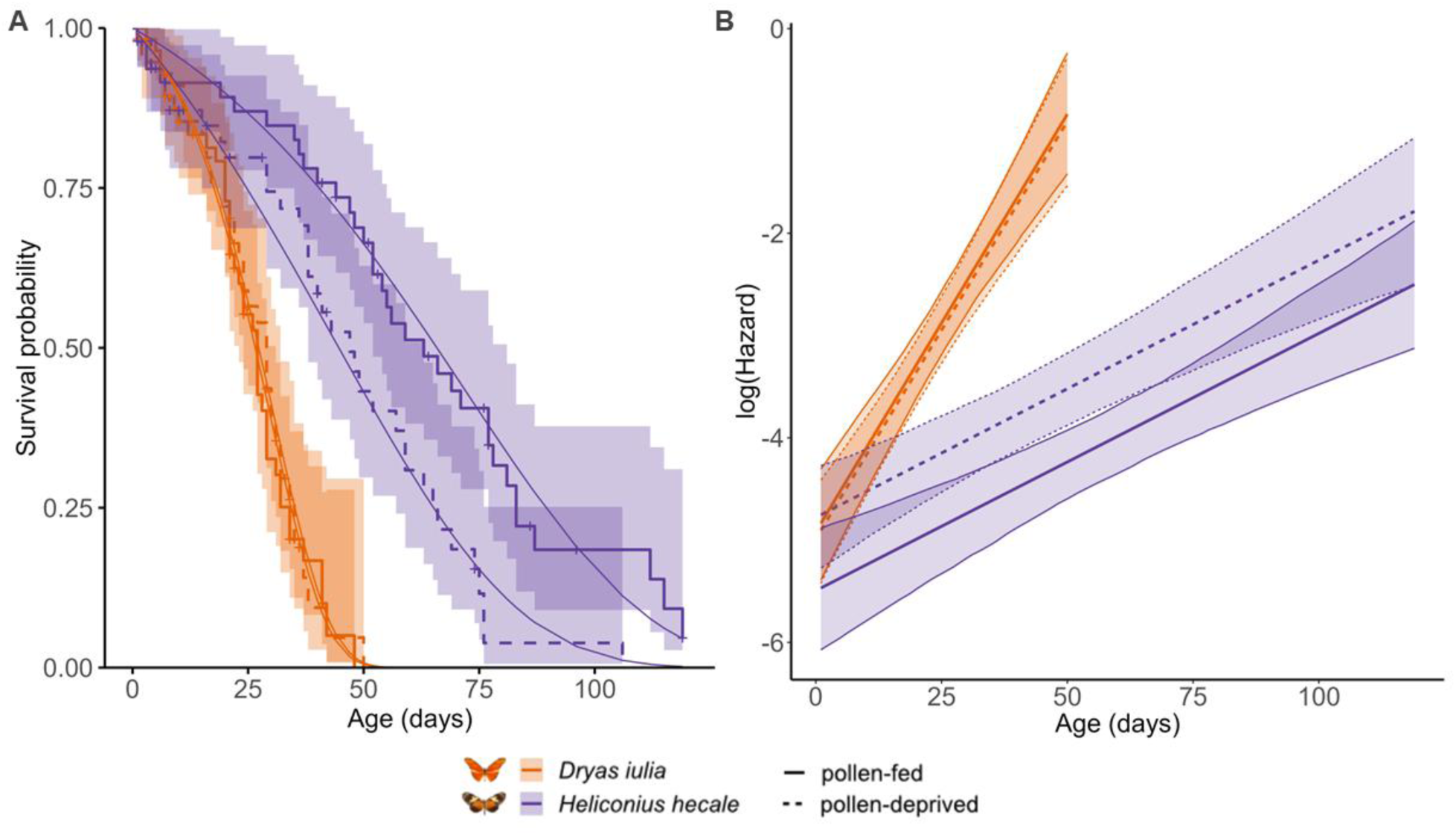
*H. hecale* shows a longer lifespan and a slowed rate of ageing compared to *D. iulia*. **A** Kaplan-Meier survival estimates and 95% confidence intervals overlaid with the corresponding parametric survival curves for the pollen-manipulation experiment cohort. “+” indicates a censored data point. **B** log(Hazard) (equivalent to log[likelihood of death]) curves and 95% confidence intervals resulting from the transformation of the parametric survival function shown in **A** to the hazard function. An increase in the intercept on this graph represents an increase in baseline mortality (α); an increase in the slope represents an increase in the rate of ageing (β).

**Table 4:** Ageing parameters for the pollen-manipulation experiment cohort.

| **Species** | **Feeding habit** | **Median lifespan (days)** | **Maximum lifespan (days)** | **Baseline mortality ($\alpha$)** | **Rate of ageing ($\beta$)** |
| --- | --- | --- | --- | --- | --- |
| *Heliconius hecale* | pollen-fed | 63 | 119 | 0.0044 | 0.025 |
|  | pollen-deprived | 47 | 106 | 0.0087 | 0.025 |
| *Dryas iulia* | pollen-fed | 27 | 48 | 0.0077 | 0.081 |
|  | pollen-deprived | 29 | 50 | 0.0071 | 0.081 |

### Pollen-deprivation specifically accelerates body mass decline in *Heliconius*

Both *H. hecale* and *D. iulia* showed an age-related decline in body mass regardless of diet treatment. However, there was a significant interaction between species, diet, and age (*F*_1, 376.62_ = 9.69, *p* = 0.002), such that pollen-deprived *H. hecale* showed a steeper body mass decline with age than the pollen-fed group, losing an estimated 3.50% of their body mass per week, compared with 1.06% for pollen-fed butterflies (Fig. 3; *F*_1, 301.09_ = 4.18, *p* < 0.001). In contrast, and in keeping with the lack of a diet effect on longevity, in *D. iulia* there was no difference in rate of body mass decline between diet treatments, with both groups losing an estimated 6.50% of their body weight per week (Fig. 3; χ^2^ = 0.28, *p* = 0.599). Although the pollen-deprived group overall weighed an estimated 9.54% less than the pollen-fed group at the onset of the experiment (*F*_1, 128.68_ = 5.08, *p* = 0.026), there was no interaction between age and pollen deprivation (χ^2^ = 0.28, *p* = 0.599). Across both species, there was a significant interaction between age and sex, with body mass in males declining more steeply with age than in females (*F*_1, 376.62_ = 9.69, *p* = 0.002), an effect which persisted across the full *H. hecale* lifespan (*F*_1, 301.41_ = 4.18, *p* = 0.042). There was also a species-specific effect of longevity, whereby longer-lived *H. hecale* individuals were generally heavier (*F*_1, 97.82_ = 8.32, *p* = 0.005), whereas this was not the case for *D. iulia* (*F*_1, 137.52_ = 0.01, *p* = 0.924).

**Fig. 3:**
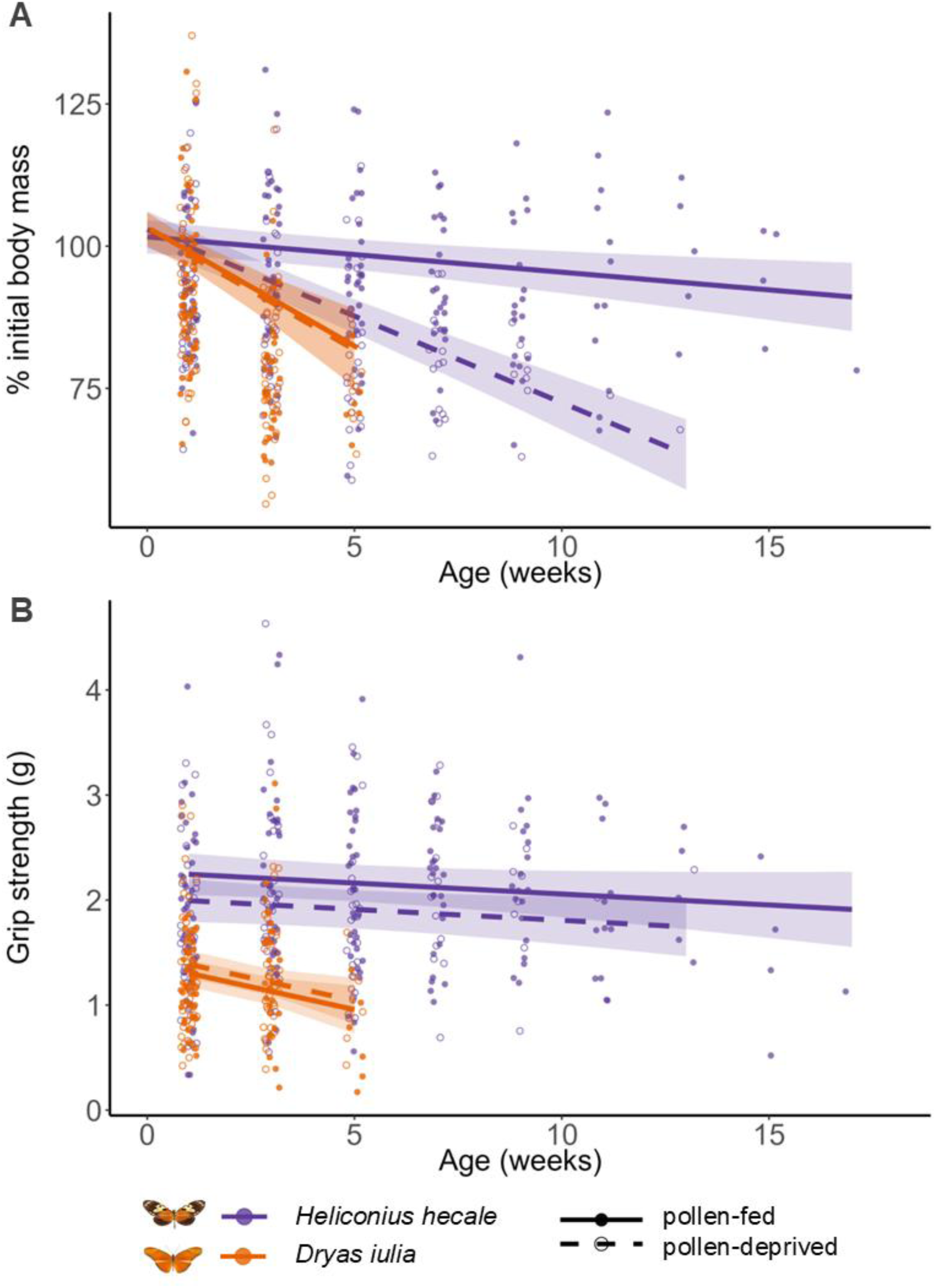
*H. hecale* shows slowed functional senescence in comparison with *D. iulia*. **A** Change in body mass with age. Both *H. hecale* and *D. iulia* show a decline in body mass with increasing age, but this is steeper in *D. iulia* and in pollen-deprived *H. hecale*. Dots represent individual data points, normalised to initial body mass to better visualise age-related change. **B** Change in grip strength with age. Grip strength declines with age in *D. iulia*, but not in *H. hecale,* although pollen-deprived *H. hecale* were weaker overall. Dots represent individual data points indicating the maximum weight a butterfly was capable of lifting with its true legs. All regression lines, with 95% confidence intervals, are from LMM analysis and show predicted traits for each species / diet combination.

### Muscle function declines with age in D. iulia, but not in H. hecale

Grip strength weakened with age in *D. iulia* (Fig. 3; *F*_1, 99.37_ = 6.78, *p* = 0.011), with butterflies pulling an estimated 0.35g less on week 5 than on than week 1, a reduction of 25.67%. However, this was not true of *H. hecale*, which did not show signs of a decline in grip strength with age, even across their much longer lifespan (Fig. 3; *F*_1, 228.82_ = 2.62, *p* = 0.107). Diet did not have an impact on grip strength in *D. iulia* (Fig. 3; *F*_1, 83.20_ = 1.14, *p* = 0.289), but in *H. hecale*, pollen-deprived butterflies were weaker than the pollen-fed group (Fig. 3; *F*_1, 72.90_ = 5.05, *p* = 0.028), pulling an estimated 0.25g less, a reduction of 12.15%. However, there was no interaction between age and diet in *H. hecale* (Fig. 3; χ^2^_1_ = 2.30, *p* = 0.130), meaning that the impact of pollen-deprivation was consistent across the full lifespan. In *H. hecale*, females were stronger than males (*F*_1, 71.65_ = 4.83, *p* = 0.031), pulling an estimated 0.23g more (12.34%), while there was no sex difference in *D. iulia* (χ^2^_1_ = 0.53, *p* = 0.467). Eclosion mass was also a significant predictor of grip strength across all butterflies (*F*_1, 180.82_ = 2.62, *p* < 0.001), with heavier butterflies capable of pulling heavier loads. Once again, there was a species-specific effect of longevity, whereby longer-lived *H. hecale* individuals were stronger (*F*_1, 97.01_ = 4.29, *p* = 0.041), while this was not the case for *D. iulia* (*F*_1, 114.93_ = 2.25, *p* = 0.136).

## Discussion

Research into long-lived species across the animal kingdom holds extraordinary potential for uncovering novel mechanisms of healthy ageing. In particular, the diversity of lifespan found in insects [9], with their experimental tractability and long history of ageing research [45], presents a fertile ground for studying the evolution of extended life. Butterflies in the *Heliconius* genus have previously been reported to live up to 6 months [15], suggesting a major lifespan extension over their non-pollen-feeding Heliconiini relatives [16, 18] and positioning them as a potential candidate system for such research. However, the paucity of published longevity data for many species within this tribe has previously limited the reliability of this lifespan extension. Collating maximum longevities from decades of published research on Heliconiini, and utilising the presence of these charismatic butterflies in commercial butterfly houses for additional data on maximum lifespan, we reveal 25-fold variation in recorded maximum lifespan across the tribe. This range far exceeds previous estimates, and is among the largest ever recorded for such closely-related taxa (with comparable differences reported only for rockfishes [46]). The maximum reported lifespan of 348 days for *Heliconius hewitsoni* also places it as the longest for any butterfly previously recorded in the scientific literature, comfortably outstripping the previous record (*Euphaedra medon*, 293 days [47]) by almost two months, and within range of *Myscelia cyaniris* (380 days), which we present as the longest-lived species to date based on data from butterfly exhibitors (Supplementary Data 2). Our data on parameters of ageing across the Heliconiini tribe, combined with a more detailed analysis of two focal longer- and shorter-lived species, show that the evolution of a novel pollen-feeding behaviour in *Heliconius* facilitated the evolution of slowed actuarial and physiological senescence and increased lifespan in this genus. Our results provide a much-needed foundation for future research into mechanisms of longevity in this new model system for the study of long life. Here we discuss three main results: i) that pollen-fed *Heliconius* outlive pollen-deprived *Heliconius*, but that ii) *Dryas iulia* gains no passive benefit of pollen; and that iii) both with and without pollen, *Heliconius* show reduced ageing over the outgroup Heliconiini, suggesting heritable mechanisms of delayed senescence.

### Pollen-feeding in *Heliconius*: direct, plastic benefits to fitness through evolved mechanisms

Results from survival analysis of *Heliconius hecale* showed direct, plastic benefits of the pollen-fed diet treatment to lifespan, with a 16-day increase in median lifespan. This is consistent with the direction of previous results in *Heliconius charithonia* [16], and also aligns with similar work showing longevity increases in pollen-fed bees [48], suggesting that pollen-feeding might provide convergent longevity benefits across these species. Parametric survival analysis shows that this increased lifespan in pollen-fed *H. hecale* is mediated through a reduction in baseline mortality independent of age. This may suggest that pollen-deprived individuals are more vulnerable to extrinsic hazards such as predation, starvation, or infectious diseases [28], which aligns with some of the potential nutritional benefits of dietary pollen. For example, pollen contains a high proportion of lipids [49], which are important in insect immunity [50], and may directly contribute to increased lifespan through enhanced immune defences.

These direct benefits of pollen consumption on physiological condition irrespective of age are neatly mirrored in the superior performance of pollen-fed *H. hecale* in the grip strength assay, which assesses muscle function. The consistency of this enhanced grip strength across the entire lifespan suggests that it is not due to beneficial impacts of pollen-derived amino acids on musculature, in which case we would expect to see an interaction between diet and age, whereby pollen-deprived butterflies gradually grew weaker as they got older. It may instead reflect a greater energy budget in these butterflies, both due to pollen-derived lipids, which are important energy storage compounds [51], and due to pollen-derived amino acids. For example, the amino acid proline plays a critical role in energy storage in insect muscle and haemolymph [19, 52], and has been found in high proportions in the pollen of *Psiguria*, the preferred pollen source of *Heliconius* used in this study [53]. Regardless, improved physiological condition may have ecological relevance for fitness in pollen-fed *Heliconius*, conferring advantages in competition for resources, or through enhanced predator avoidance, as reflected in their reduced baseline mortality. None of these direct plastic fitness benefits of pollen was found in *D. iulia*, despite the possibility of incidental amino acid uptake given the capacity of pollen to release free amino acids in nectar [19]. This suggests that *Heliconius* have not only evolved the ability to actively collect and digest pollen, but also the physiological adaptations necessary to fully exploit its nutritional benefits – adaptations that are absent in *D. iulia* and likely in the other non-pollen-feeding Heliconiini.

### Indirect benefits of pollen: resource allocation and the disposable soma theory

While previous work has suggested that *Heliconius’* lifespan extension is an exclusively plastic response to adult pollen intake, reporting that pollen-deprivation reduces *Heliconius* lifespan to that of their non-pollen-feeding Heliconiini relatives [16], our results demonstrate a 20-day increase in median lifespan from *D. iulia* to pollen-deprived *H. hecale.* Life history theory predicts that increased investment in lifespan should come with concomitant costs to other traits, such as fecundity [54]. The disposable soma theory similarly places the evolution of ageing within a resource allocation framework [55], arguing that it is the compromise between allocation of limited resources to either reproduction or somatic maintenance that ultimately leads to senescence [56]. However, the boundaries of this framework may not be so tightly fixed if there is a change in the reserve of available resources. Both on a plastic, individual level [57], and on an evolutionary timescale [9, 58, 59], enhanced nutrition can increase an organism’s overall fitness by positively impacting one life history trait without incurring a complementary cost. The evolution of pollen-feeding in *Heliconius* seems to have loosened life history constraints by adding a valuable additional nutritional resource, shifting the burden of reproduction to the adult stage [16].

Support for this shift comes from isotopic evidence for transfer of essential amino acids from pollen to eggs [23], as well as the finding that while larval nutrition and development time are similar across pollen-feeding and non-pollen-feeding Heliconiini [60], the proportion of nitrogenous resources allocated to reproduction at eclosion is inversely correlated to expected adult nitrogen intake [61]. This suggests that in *Heliconius,* a greater proportion of larval-derived resources may instead be allocated to somatic maintenance and longevity, explaining the lifespan increase in pollen-deprived *H. hecale*. This is supported by the results of our grip strength assay, in which both pollen-fed and pollen-deprived *H. hecale* lack senescence in physiological condition, in contrast to ageing *D. iulia*, making it unlikely that the burden of somatic maintenance is shifted to adult-acquired nitrogenous reserves. This is further reinforced by the observed patterns of body mass decline, which is thought to result from the depletion of stored reproductive reserves [20, 31], and is attenuated in *H. hecale* as compared to *D. iulia* in accordance with their enhanced adult diet. Without the expected supplementation of nitrogenous reserves, pollen-deprived *H. hecale* suffer a steeper decline in body mass as reproductive reserves rapidly deplete. However, this decline is still intermediate between that of *D. iulia* and the pollen-fed *H. hecale*. Tied with the increased lifespan and lack of physiological senescence shown in pollen-deprived *H. hecale*, this suggests some larval reserves are kept aside for somatic maintenance regardless of reproductive deficits imposed by a nitrogen-poor adult diet.

### Evolution of long life and slowed ageing in *Heliconius*

Placing these findings in an evolutionary context suggests that the transition to pollen-feeding in *Heliconius*, and the prolonged reproductive lifespan it permits [16], caused the “selection shadow” central to evolutionary theories of ageing to retreat to higher reaches of the lifespan. Increased reproductive longevity exposes older ages to selection, favouring the evolution of pro-longevity mechanisms; for example, the proposed shift in resource allocation towards somatic maintenance suggested by our results in *H. hecale* (Supplementary Note 9). Depending on the timing of this dietary innovation, such mechanisms may have evolved with varying degrees of efficacy in different *Heliconius* lineages, possibly related to species-specific differences in ecology. This could explain the wide variation in lifespan between *Heliconius* species, with maximum lifespans for some even falling within the range of the outgroup Heliconiini genera (Fig. 1, Table 1). It is possible that some of this variation may reflect differences in response to caged environments. However, a major strength of the data presented here is its combination of records from semi-natural butterfly house exhibits and mark-release recapture experiments in the wild in addition to those from caged insectary populations (Supplementary Data 1), meaning our inferences are robust to any differences between species in response to any one experimental set-up. Our collation of multiple records per species across many studies (Tables 1–4, Supplementary Data 1) also highlights the limitations of using maximum lifespan as the sole metric of ageing, as large intra-species differences suggest that these values are often driven by single long-lived individuals.

Beyond maximum lifespan, other, more informative metrics of ageing across the Heliconiini tribe also support the idea that *Heliconius’* lifespan extension was facilitated by the evolution of pollen-feeding. Survival data from 17 Heliconiini species also show differences in median lifespan, baseline mortality (𝛼), and rate of ageing (𝛽) across the tribe, as well as within the *Heliconius* genus. The transition to pollen-feeding corresponds to reduced baseline mortality (𝛼) and rate of ageing (𝛽), and in accordance with this, lengthened lifespans in *Heliconius*. Although we lack longevity data from the Aoede clade, the only non-pollen-feeding *Heliconius* (previously considered a separate genus, *Neruda*) [17], particular support for an association with pollen-feeding and/or other derived *Heliconius* traits comes from our data from the non-pollen-feeding *Eueides*, as the genus most closely related to *Heliconius* [17]. Despite this closer phylogenetic proximity, the maximum reported lifespan for *Eueides isabella* of 50 days (from our own data in Table 3) places it comfortably within the range of maximum lifespans displayed by the other non-pollen-feeding Heliconiini outgroups, and below that of all *Heliconius* data. While our data reflects a single evolutionary transition at the base of *Heliconius*, the overlapping habitats, host-plants, and common juvenile life histories across Heliconiini leaves pollen-feeding as the only clear ecological difference that distinguishes *Heliconius* from outgroup genera and can explain the observed shift in adult lifespan.

This phylogenetically-broad survival analysis provided the required background to then narrow our focus to our two representative longer- and shorter-lived species, *H. hecale* and *D. iulia*, to better understand the underlying basis of *Heliconius*’ longevity. While lifespan data is informative, modelling age-specific mortality via parametric survival analysis can provide additional information on patterns of ageing, allowing for deeper insights into the biological underpinnings of such lifespan extensions [25]. An “extended lifespan” does not necessarily mean that senescence has been delayed, but may instead reflect a reduction in baseline mortality that results in greater survival irrespective of age, as has been shown for example across mammalian females [62]. The results of the pollen-manipulation experiment show that while *H. hecale* and *D. iulia* show similar baseline mortality, *H. hecale* has evolved a slower rate of ageing, resulting in the observed lifespan extension. This slowed rate is maintained even under pollen-deprivation, suggesting that this parameter is unrelated to the short-term plastic effects of adult diet (Fig. 2). Our results are of particular significance considering *D. iulia*’s position as one of the longest-lived non-pollen-feeding outgroups (Tables 1-3), as we find clear differences in metrics of ageing in our experiments despite the apparent capacity for lengthy lifespans in some individuals of the species. While the wide variation in both of these parameters across the tribe makes it unlikely that this pattern is precisely generalisable to all Heliconiini, our combined results across several cohorts consistently suggest a slowed rate of ageing as a major contributor to lengthened lifespan in *Heliconius*.

A reduction in the rate of ageing implies the presence of mechanisms that delay or slow physiological senescence [28]. This is illuminated by results from the grip strength assay, which show an age-related decline in grip strength in *D. iulia* and provide for the first time a reliable index of physiological senescence in Heliconiini butterflies. This assay has been proposed to measure whole organism performance in butterflies [34], citing similar metrics that predict fitness in beetles [35], and hand grip strength is also used as a biomarker of health in ageing humans [36]. In *D. iulia*, declining grip strength is likely an indicator of deteriorating muscular health, with age-related declines in muscular structure and function also reported in several other insects [33]. This decline in *D. iulia* provides a possible physiological basis for their faster actuarial senescence as compared to *H. hecale*, and this assay links actuarial and physiological senescence in this system, with a lower overall grip strength also associated with higher baseline mortality in pollen-deprived *H. hecale*. In contrast with *D. iulia*, ageing *H. hecale* showed no sign of a similar decline in grip strength, maintaining performance even at very old ages. This shows that long past the limits of *D. iulia’s* lifespan, this deterioration is mitigated in *H. hecale*. Our findings show that in accordance with their lengthened lifespan and slowed actuarial senescence, these butterflies have also evolved a delayed physiological senescence.

## Conclusion

Our data provide the first longitudinal study of functional senescence in a long-lived butterfly genus, and present evidence for an evolved lifespan extension, slowed actuarial senescence, and delayed physiological senescence in *H. hecale*, independent of any short-term, plastic benefits of adult pollen intake. The applicability of these findings to other *Heliconius* species may depend on ecological factors such as the degree of pollen reliance [63], and would be illuminated by further studies of senescence in other *Heliconius* species. However, our data spans multiple metrics of ageing and combines several independent sources to show that any intra-*Heliconius* variation is surpassed by the broader differences between *Heliconius* and the other Heliconiini (Fig. 1). This makes our in-depth findings in *H. hecale* a productive representation of this genus, and establishes *Heliconius* as a valuable new model system for the study of extended longevity. Future work could include mechanistic investigations to explore the proximate basis for this extended “health span”, and allow us to develop a deeper understanding of the tools these butterflies are using to unpick the constraints imposed by the ageing paradigm.

## Acknowledgements

We are very grateful to the Ministerio del Ambiente in Panama for permission to collect and export samples, and to the Smithsonian Tropical Research Institute for providing the facilities to carry out much of this research. We would like to thank Laura Hebberecht-Lopez, Lina Melo, and Oscar Paneso for their assistance with rearing at the insectaries. We would also like to thank Michael Hearn and Andrew Quitmeyer for their assistance in developing *The Pullinator*, and the members of the International Association of Butterfly Exhibitors and Suppliers who provided additional maximum longevity records for Heliconiini species in their exhibits. This work was supported by a GW4 BioMed MRC DTP (J.F.), a NERC Independent Research Fellowship (NE/N014936/1) and ERC Starter Grant (758508) (S.H.M.), and funding from the US National Science Foundation (IOS 2110532) and the Smithsonian Tropical Research Institute (W.O.M.).

## References

1. Austad, S.N., Methusaleh’s Zoo: How Nature provides us with Clues for Extending Human Health Span. Journal of Comparative Pathology, 2010. 142: p. S10–S21.

2. Aristotle, *On Length and Shortness of Life*, in Parva Naturalia, W.D. Ross, Editor. 1955, Oxford University Press: Oxford.

3. Jones, O.R., et al., Diversity of ageing across the tree of life. Nature, 2014. 505(7482): p. 169–173.

4. Medawar, P.B., An Unsolved Problem of Biology. 1952, London: H.K. Lewis & Co.

5. Haldane, J.B.S., New Paths in Genetics. 1941, London: Allen & Unwin. 206 p.

6. Hamilton, W.D., The moulding of senescence by natural selection. Journal of Theoretical Biology, 1966. 12(1): p. 12–45.

7. Williams, G.C., Pleiotropy, Natural Selection, and the Evolution of Senescence. Evolution, 1957. 11(4): p. 398.

8. Fontana, L., L. Partridge, and V.D. Longo, Extending Healthy Life Span - From Yeast to Humans. Science, 2010. 328(5976): p. 321–326.

9. Carey, J.R., Insect Biodemography. Annual Review of Entomology, 2001. 46(1): p. 79–110.

10. Clancy, D.J., *Extension of Life-Span by Loss of CHICO, a* Drosophila *Insulin Receptor Substrate Protein*. Science, 2001. 292(5514): p. 104–106.

11. Tatar, M., *A Mutant* Drosophila *Insulin Receptor Homolog That Extends Life-Span and Impairs Neuroendocrine Function*. Science, 2001. 292(5514): p. 107–110.

12. Elsner, D., K. Meusemann, and J. Korb, Longevity and transposon defense, the case of termite reproductives. Proceedings of the National Academy of Sciences, 2018. 115(21): p. 5504.

13. Lucas, E.R., E. Privman, and L. Keller, Higher expression of somatic repair genes in long-lived ant queens than workers. Aging, 2016. 8(9): p. 1940–1951.

14. Beck, J. and K. Fiedler, Adult life spans of butterflies (Lepidoptera: Papilionoidea + Hesperioidea): broadscale contingencies with adult and larval traits in multi-species comparisons. Biological Journal of the Linnean Society, 2009. 96(1): p. 166–184.

15. Ehrlich, P.R. and L.E. Gilbert, *Population Structure and Dynamics of the Tropical Butterfly* Heliconius ethilla. Biotropica, 1973. 5(2): p. 69–82.

16. Dunlap-Pianka, H., C.L. Boggs, and L.E. Gilbert, Ovarian Dynamics in Heliconiine Butterflies: Programmed Senescence versus Eternal Youth. Science, 1977. 197(4302): p. 487–490.

17. Cicconardi, F., et al., Evolutionary dynamics of genome size and content during the adaptive radiation of Heliconiini butterflies. Nature Communications, 2023. 14(1).

18. Young, F.J. and S.H. Montgomery, *Pollen feeding in* Heliconius *butterflies: the singular evolution of an adaptive suite*. Proceedings of the Royal Society B: Biological Sciences, 2020. 287(1938): p. 20201304.

19. Gilbert, L.E., *Pollen feeding and reproductive biology of* heliconius *butterflies*. Proc Natl Acad Sci U S A, 1972. 69(6): p. 1403–7.

20. Boggs, C.L., Understanding insect life histories and senescence through a resource allocation lens. Functional Ecology, 2009. 23(1): p. 27–37.

21. Couto, A., et al., Rapid expansion and visual specialisation of learning and memory centres in the brains of Heliconiini butterflies. Nature Communications, 2023. 14(1).

22. Young, F.J., et al., Enhanced long-term memory and increased mushroom body plasticity in Heliconius butterflies. iScience, 2024: p. 108949.

23. O’Brien, D.M., C.L. Boggs, and M.L. Fogel, *Pollen feeding in the butterfly* Heliconius charitonia*: isotopic evidence for essential amino acid transfer from pollen to eggs*. Proceedings of the Royal Society of London. Series B: Biological Sciences, 2003. 270(1533): p. 2631–2636.

24. Pinheiro De Castro, E.C., et al., Pollen-feeding delays reproductive senescence and maintains toxicity of Heliconius erato. Peer Community Journal, 2025. 5.

25. Bronikowski, A. and T. Flatt, Aging and its demographic measurement. Nat Educ Knowl, 2010. 1.

26. Ronget, V. and J.M. Gaillard, Assessing ageing patterns for comparative analyses of mortality curves: Going beyond the use of maximum longevity. Functional Ecology, 2020. 34(1): p. 65–75.

27. Moorad, J.A., et al., A comparative assessment of univariate longevity measures using zoological animal records. Aging Cell, 2012. 11(6): p. 940–948.

28. Ricklefs, R.E. and A. Scheuerlein, Biological Implications of the Weibull and Gompertz Models of Aging. The Journals of Gerontology Series A: Biological Sciences and Medical Sciences, 2002. 57(2): p. B69–B76.

29. Brewster, A.L.E. and G.W. Otis, A Protocol for Evaluating Cost-Effectiveness of Butterflies in Live Exhibits. Journal of Economic Entomology, 2009. 102(1): p. 105–114.

30. Brown, K., *The biology of* Heliconius *and related genera*. Annual review of entomology, 1981. 26(1): p. 427–457.

31. Boggs, C.L., REPRODUCTIVE ALLOCATION FROM RESERVES AND INCOME IN BUTTERFLY SPECIES WITH DIFFERING ADULT DIETS. Ecology, 1997. 78(1): p. 181–191.

32. Pásztor, K., et al., Phenotypic senescence in a natural insect population. Ecology and Evolution, 2022. 12(12).

33. Sohal, R.S., 18 - Aging in Insects, in Biochemistry, G.A. Kerkut and L.I. Gilbert, Editors. 1985, Pergamon: Amsterdam. p. 595–631.

34. Davis, A.K., F.M. Smith, and A.M. Ballew, A poor substitute for the real thing: captive-reared monarch butterflies are weaker, paler and have less elongated wings than wild migrants. Biology Letters, 2020. 16(4): p. 20190922.

35. Lailvaux, S.P., et al., *Horn size predicts physical performance in the beetle* Euoniticellus intermedius *(Coleoptera: Scarabaeidae)*. Functional Ecology, 2005. 19(4): p. 632–639.

36. Bohannon, R.W., Grip Strength: An Indispensable Biomarker For Older Adults. Clinical Interventions in Aging, 2019. **Volume** 14: p. 1681–1691.

37. R Core Team, R: A Language and Environment for Statistical Computing. 2023, R Foundation for Statistical Computing: Vienna, Austria.

38. Therneau, T.M., A Package for Survival Analysis in R. 2023.

39. Therneau, T.M., coxme: Mixed Effects Cox Models. 2022.

40. Jackson, C., *flexsurv: A Platform for Parametric Survival Modeling in R*. Journal of Statistical Software, 2016. 70(8): p. 1–33.

41. Gompertz, B., On the Nature of the Function Expressive of the Law of Human Mortality, and on a New Mode of Determining the Value of Life Contingencies. Philosophical Transactions of the Royal Society of London, 1825. 115: p. 513–583.

42. Colchero, F., O.R. Jones, and M. Rebke, BaSTA: an R package for Bayesian estimation of age-specific survival from incomplete mark-recapture/recovery data with covariates. Methods in Ecology and Evolution, 2012. 3(3): p. 466–470.

43. Revell, L.J., *phytools 2.0: an updated R ecosystem for phylogenetic comparative methods (and other things)*. PeerJ, 2024. 12: p. e16505.

44. Bates, D., et al., Fitting Linear Mixed-Effects Models Using lme4. Journal of Statistical Software, 2015. 67(1): p. 1–48.

45. Promislow, D.E.L., T. Flatt, and R. Bonduriansky, The Biology of Aging in Insects: From Drosophila to Other Insects and Back. Annual Review of Entomology, 2022. 67(1): p. 83–103.

46. Kolora, S.R.R., et al., Origins and evolution of extreme life span in Pacific Ocean rockfishes. Science, 2021. 374(6569): p. 842–847.

47. Molleman, F., et al., Extraordinary long life spans in fruit-feeding butterflies can provide window on evolution of life span and aging. Experimental Gerontology, 2007. 42(6): p. 472–482.

48. Schmidt, J.O., S.C. Thoenes, and M.D. Levin, *Survival of Honey Bees*, Apis mellifera *(Hymenoptera: Apidae), Fed Various Pollen Sources*. Annals of the Entomological Society of America, 1987. 80(2): p. 176–183.

49. Roulston, T.H. and J.H. Cane, Pollen nutritional content and digestibility for animals. Plant Systematics and Evolution, 2000. 222(1): p. 187–209.

50. Wrońska, A.K., et al., Lipids as a key element of insect defense systems. Frontiers in Genetics, 2023. 14.

51. Arrese, E.L. and J.L. Soulages, Insect Fat Body: Energy, Metabolism, and Regulation. Annual Review of Entomology, 2010. 55(1): p. 207–225.

52. Teulier, L., et al., Proline as a fuel for insect flight: enhancing carbohydrate oxidation in hymenopterans. Proceedings of the Royal Society B: Biological Sciences, 2016. 283(1834): p. 20160333.

53. Cardoso, M.Z. and L.E. Gilbert, Pollen feeding, resource allocation and the evolution of chemical defence in passion vine butterflies. Journal of Evolutionary Biology, 2013. 26(6): p. 1254–1260.

54. Stearns, S.C., The evolution of life histories. 1992, Oxford ;: Oxford University Press.

55. Kirkwood, T.B.L., Evolution of ageing. Nature, 1977. 270(5635): p. 301–304.

56. Jones, O.R., et al., The Disposable Soma Theory: Origins and Evolution, in The Evolution of Senescence in the Tree of Life. 2017, Cambridge University Press: Cambridge. p. 23–39.

57. Nylin, S. and K. Gotthard, Plasticity in Life-History Traits. Annual Review of Entomology, 1998. 43(1): p. 63–83.

58. Kaplan, H., et al., *A theory of human life history evolution: Diet, intelligence, and longevity.* Evolutionary Anthropology: Issues, News, and Reviews, 2000. 9(4): p. 156–185.

59. Swanson, E.M., et al., Nutrition shapes life-history evolution across species. Proceedings of the Royal Society B: Biological Sciences, 2016. 283(1834): p. 20152764.

60. Hebberecht, L., et al., *The evolution of adult pollen feeding did not alter postembryonic growth in* Heliconius *butterflies*. Ecology and Evolution, 2022. 12(6).

61. Boggs, C.L., Nutritional and Life-History Determinants of Resource Allocation in Holometabolous Insects. The American Naturalist, 1981. 117(5): p. 692–709.

62. Lemaître, J.-F., et al., Sex differences in adult lifespan and aging rates of mortality across wild mammals. Proceedings of the National Academy of Sciences, 2020. 117(15): p. 8546–8553.

63. Boggs, C.L., J.T. Smiley, and L.E. Gilbert, *Patterns of pollen exploitation by* Heliconius *butterflies*. Oecologia, 1981. 48(2): p. 284–289.

64. Seixas, R.R., et al., *Population Biology of the Sand Forest Specialist Butterfly* Heliconius *hermathena hermathena (Hewitson) (Nymphalidae: Heliconiinae) in Central Amazonia).* The Journal of the Lepidopterists’ Society, 2017. 71(3): p. 133–140.

65. Kelson, R. Searching for Methuselah: Butterfly Longevity Revisited. in Invertebrates in Captivity Conference. 2008. Sonoran Arthropod Studies Institute, Tucson, AZ.

66. Brown, K.S., The heliconians of Brazil (Lepidoptera: Nymphalidae). Part III. Ecology and biology of Heliconius nattereri, a key primitive species neat extinction, and comments on the evolutionary development of Heloconius and Eueides. Zoologica : scientific contributions of the New York Zoological Society., 1972. 57(2): p. 41–69.

67. Mallet, J.L.B. and D.A. Jackson, *The ecology and social behaviour of the Neotropical butterfly* Heliconius xanthocles *Bates in Colombia*. Zoological Journal of the Linnean Society, 1980. 70(1): p. 1–13.

68. Menacé Almea, M.A., C. Belezaca, and M.A. Lara Valarezo, *Análisis en condiciones semicontroladas de la biología del gusano defoliador (*Dione juno juno*) de la maracuyá (passiflora edulis), en el litoral del Ecuador*. Revista Universidad y Sociedad, 2019. 11: p. 215–219.

