## Supplementary Materials for "Evolution of increased longevity and slowed ageing in a genus of tropical butterfly"

### 1. Supplementary Methods

##### *1.1 Heliconiini maximum reported lifespan data collation*

Data on maximum reported lifespan for species across the Heliconiini tribe (Table 1, Supplementary Data 1) was first collated from a literature search within PubMed, Google Scholar, and Web of Science for long-term studies of *Heliconius* and other Heliconiini butterflies, using combinations of the search terms "*Heliconius*", "Heliconiini", "lifespan", "longevity", "mark", "recapture", and "population dynamics", between January and April 2021. However, considering the popularity of Heliconiini species in commercial butterfly houses, such exhibits were also contacted to enquire if they had collected lifespan data, as has been done previously [1]. A list of active members of the International Association of Butterfly Exhibitors and Suppliers was obtained and members were systematically contacted via email. One study in particular (Kelson, unpublished) was responsible for many of the highest maximum longevity records; for further details of this study design, see Supplementary Note 7.

##### *1.2 Butterfly husbandry and survival data collection*

All Panamanian butterflies were collected under permit number SE/A-82-19 and SE/A-14-18.

*1.2.1 Multi-species cognitive experiment cohort*

To move beyond maximum lifespan and better understand patterns of ageing across Heliconiini, survival data from [2] was taken for *Heliconius hecale melicerta, Heliconius melpomene rosina*, *Dryadula phaetusa*, and *Agraulis vanillae*. Survival data for *Dryas iulia* was taken from a similar cognitive dataset (Foley *et al*., in prep) generated under the same experimental conditions. In both studies, all butterflies were reared from stock populations at the Smithsonian Tropical Research Institute (STRI) outdoor insectaries in Gamboa, Panama. These populations were established from wild-caught individuals collected within a 2km radius of Gamboa, between January-May 2019 (for *H. melpomene*, *H. hecale*, *D. phaetusa*, and *A. vanillae*), or between January-November 2022 (for *D. iulia*). Eggs were collected daily from host-plants within these stock cages, and hatched larvae were reared in mesh pop-up cages and fed *ad libitum* on leaves of their preferred *Passiflora* host-plants. *H. melpomene* was fed on *P. triloba*, *H. hecale* was fed on *P. vitifolia*, and *D. phaetusa*, *A. vanillae*, and *D. iulia* were fed on *P. biflora*.

Upon eclosion, butterflies were marked with a unique ID and placed into 2m (L) x 3m (W) x 2m (H) cages containing a rack of 24 artificial feeders arranged in a 4 x 6 grid and placed centrally in the cage. Feeders contained approximately 0.5ml of a sucrose-protein solution (25% w/v sucrose and 5% w/v Vetark Critical Care Formula), replaced daily. A single non-flowering *Palicourea elata* plant was also placed in each cage as a roosting site. Adult butterflies were subjected to long-term memory assays based on the association of a colour with a food reward, which imposed regular bouts of food deprivation during testing periods as well as frequent handling. Further details of these assays may be found in [2].

Upon completing the cognitive experiment, butterflies were maintained until the end of their natural lifespans. Cages were checked daily for dead individuals, death dates were recorded, and missing or predated individuals were censored at the age at which they were last seen alive. In total, survival data from both sexes was analysed for 175 individuals of *A. vanillae* (12 censored observations), 263 individuals of *D. iulia* (38 censored observations), 108 individuals of *D. phaetusa* (10 censored observations), 120 individuals of *H. hecale* (56 censored observations), and 103 individuals of *H. melpomene* (31 censored observations).

*1.2.2 Semi-natural “mark-release-recapture” cohort*

To gather survival data across a wider sampling of Heliconiini, butterflies of 20 different species were released in a semi-natural enclosure located at the STRI insectaries. As wide a range of species across the phylogeny as possible was included, representing a range of different ecological niches; therefore, provenance of these stock populations varied depending on the species, with many established from local collecting trips near Gamboa or elsewhere in Panama, and others established using pupae from butterfly farms from other countries in the region. All individuals were reared from stock populations at the STRI insectaries, between January and November 2022.

Larvae of each species were reared on leaves of their preferred *Passiflora* host-plants. Upon eclosion, adults were marked with a unique ID and released into a large 11m (L) x 11m (W) x 6m (H) cage similar to those found in commercial butterfly houses. This cage was designed to mimic the natural habitat of these butterflies as closely as possible, with a wide variety of trees, *Passiflora* host-plants, and other flowering plants creating a semi-natural environment for the butterflies inhabiting it. Floral nectar sources were also supplemented by artificial feeders containing a 20% w/v sucrose solution, replaced every 2 days.

In total, 959 butterflies of 20 different Heliconiini species were released into the cage, including: *A. vanillae* (n = 12), *D. iulia* (n = 34), *Dione juno* (n = 45), *D. phaetusa* (n = 30),

*Eueides isabella* (n = 59), *Heliconius atthis* (n = 33), *Heliconius charithonia* (n = 3), *Heliconius cydno* (n = 19), *Heliconius doris* (n = 32), *Heliconius erato* (n = 46), *Heliconius hewitsoni* (n = 15), *H. hecale* (n = 47), *Heliconius himera* (n = 2), *Heliconius ismenius* (n = 20), *H. melpomene* (n = 285), *Heliconius numata* (n = 94), *Heliconius pachinus* (n = 19), *Heliconius sapho* (n = 81), *Heliconius sara* (n = 70), and *Philaethria dido* (n = 13). Species selected were those being reared at the STRI insectaries at the time of the experiment. All *Heliconius* included in this study are pollen-feeding species, and data for the four non-pollen feeding *Heliconius* in the Aoede clade are unfortunately not available. This reflects the understudied nature of these species, which occur at low densities, have a derived host plant, and are therefore challenging to work with [3]. Sample number reflects availability of pupae. Both sexes were represented for all species.

Data collection was carried out on a weekly basis, with some omissions due to time constraints. Approximately 15 minutes were spent patrolling the cage and recording the IDs of any butterflies visually identified and alive on that date. To ensure minimal intervention, butterflies were not recaptured or handled. Cage checks were often conducted mid-morning, when Heliconiini butterflies are at their busiest, but an effort was made to also check at other times to re-sight individuals with different active periods.

Of the 959 butterflies released into the cage, 448 individuals were re-sighted on at least one occasion. Due to the size of the cage and semi-natural conditions, dead individuals were rarely recovered before being eaten by ants, or otherwise deteriorated. In the event that a dead individual's body was recovered intact, it was assumed to have died that day, and so its death date was noted.

##### *1.2.3 Pollen-manipulation experiment cohort*

All butterflies used in the longitudinal pollen-manipulation experiments were reared from stock populations at the STRI outdoor insectaries in Gamboa, Panama, between January and November 2022. Stock populations of *H. hecale* and *D. iulia*, selected for their local abundance and ease of rearing, were established from wild-caught individuals collected within a 2km radius of Gamboa. New individuals were collected and added to stock populations approximately bi-weekly to ensure genetic diversity, with stock populations of each species in total comprising of approximately 60 females and 40 males across the 10-month field season. Stocks were maintained in 2m (L) x 2m (W) x 2m (H) cages containing artificial feeders filled with a 20% w/v sucrose and 10% w/v organic, pesticide-free bee pollen solution, changed every 2 days. All stock cages contained flowering *Palicourea,*

*Lantana,* and *Stachytarpheta* plants, and *H. hecale* stocks were also provided with fresh *Psiguria* flowers daily.

Eggs were collected daily from host-plants within these stock cages. Hatched larvae were reared in pop-up mesh cages and fed *ad libitum* on shoots of one of their preferred *Passiflora* host-plants, depending on availability: *P. biflora*, *P. auriculata, P. pittieri,* or *P. edulis* for *D. iulia*; and *P. nitida, P. riparia,* or *P. vitifolia* for *H. hecale*.

Upon eclosion, butterflies were sexed and weighed, their forewings were measured, and they were marked with a unique ID. They were then randomly assigned to either a pollen-fed or pollen-deprived treatment, placed in the corresponding cage, and taught to feed on the artificial feeders by extending their probosces into the sucrose solution. Pollen-fed cages contained approximately 12 flowering *Palicourea*, *Lantana,* and *Stachytarpheta* plants, as well as fresh *Psiguria* flowers, replaced daily. Pollen-fed cages also contained 4 central artificial feeders filled with a 20% w/v sucrose and 10% w/v organic bee pollen solution, changed daily. Pollen-deprived cages contained approximately 12 non-flowering *Palicourea*, *Lantana,* and *Stachytarpheta* plants, as well as 4 central artificial feeders and 20 red star-shaped feeders hung around the plant foliage, all filled with a 20% w/v sucrose solution, also changed daily. This was to ensure feeding opportunities were as standardised as possible between cages. All cages measured approximately 4m (L) x 3m (W) x 2m (H), and contained *P. biflora* and *P. vitifolia* host-plants. Cages were checked daily for dead individuals, death dates were recorded, and missing or predated individuals were censored at the age at which they were last seen alive. In total, survival data from both sexes was collected across the lifespan of 96 individuals of *H. hecale* (47 pollen-fed, 49 pollen-deprived; 26 censored observations) and 116 individuals of *D. iulia* (57 pollen-fed, 57 pollen-deprived; 36 censored observations).

##### *1.3 Functional senescence assays*

Butterflies from the pollen-manipulation experiment cohort ($n_{H.\ hecale}$ = 96; $n_{D.\ iulia}$ = 116) were assayed with a battery of tests every two weeks, beginning one week after eclosion, and continuing until their natural death (Fig. S1). During testing, individuals were removed from their cages at approximately 10:00, and subjected first to the flight behaviour assay, followed by the body mass and grip strength assays (see below). The flight behaviour assays were performed in the outdoor insectaries, and the body mass and grip strength assays were performed in the neighbouring laboratory. These assays were chosen to assess various indices of physiological senescence in these butterflies, as several insects show age-related declines in flight capacity [4-6], body mass [7, 8], and muscle function [5]. Testing generally concluded by 13:00, after which butterflies were returned to their respective cages.

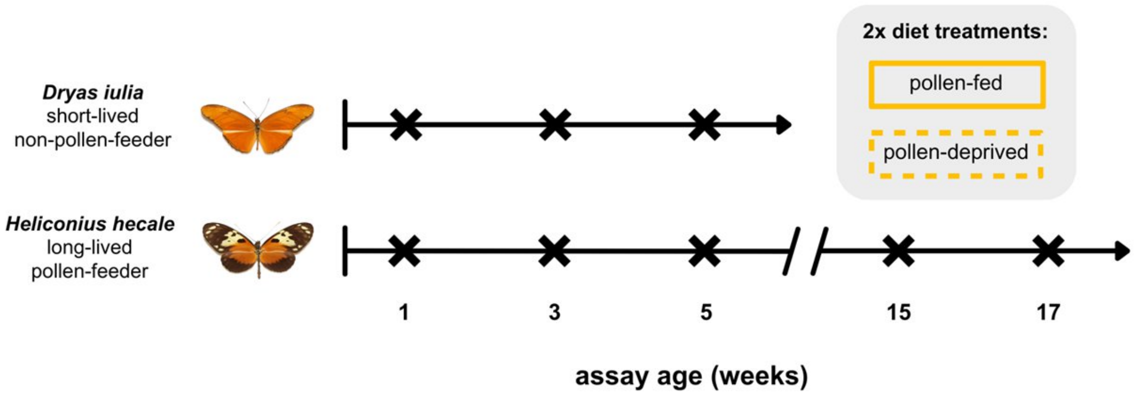

**Fig. S1: Experimental design.**

Individuals of both *H. hecale* and *D. iulia* were maintained under either a pollen-fed or pollen-deprived treatment and assayed on a biweekly basis until death, beginning at day 7 post-eclosion and continuing every 2 weeks thereafter.

##### *1.3.1 Body mass and grip strength assays*

Butterflies were first weighed using a Sartorius Entris balance with 1mg resolution. Grip strength was then assayed using a method adapted from [9] (Fig. S2). A custom-built device, denoted *The Pullinator*, was placed on to the balance, which was then tared. *The Pullinator* consisted of a lightweight wooden base upon which was mounted a small wooden perch of diameter 9.5mm, wrapped in 400 grit sandpaper to standardise friction. Butterflies were held by their wings and lowered on to the perch until they grasped it with their true legs. They were then gently pulled upwards until they released the perch, and the peak negative reading on the balance from this exercise was taken as a measure of grip strength. This was repeated five times for each individual, and the maximum reading was used for statistical analysis.

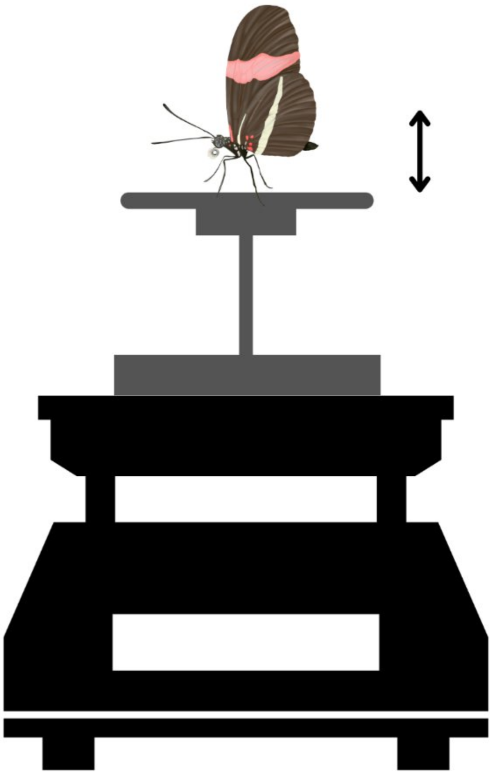

**Fig. S2:** ***The Pullinator*** **grip strength assay apparatus.**

*1.3.2 Flight behaviour assay*

Each individual was released from a height of 1.5m at the end of a narrow flight arena measuring 13m (L) x 0.75m (W) x 2m (H). Individuals were monitored for a 45-second interval, and total time spent in active flight was recorded. All butterflies were assayed twice, with an interval of at least 5 minutes between each assay. Measurements taken from the assay with the maximum time spent in active flight were then used for statistical analysis. Light intensity (illuminance) within the flight arena was recorded before each assay using a digital light meter to account for any impact of weather on flight behaviour. Daily weather variables including the ultraviolet index (UVI) and whether it was raining were also noted at the time of experiments. Results from this assay are presented in Supplementary Note 1.

##### *1.4 Statistical analyses*

All statistical analyses were conducted using R v.4.3.1 [10]. For all individuals, “age” was measured beginning from time of adult eclosion, and so does not reflect the larval and pupal stages. These generally last about 3 weeks in Heliconiini butterflies, with only minor differences (1-2 days) between species [11]. All survival analyses presented in this thesis are therefore conducted using only adult lifespan, coinciding with the earliest age at reproduction as is traditionally suggested when modelling senescence [12, 13].

##### *1.4.1 Non-parametric and semi-parametric survival analyses*

The package *survival* v3.5-5 [14] was used to generate Kaplan-Meier survival curves and corresponding median lifespan for all cohorts with the exception of the semi-natural "mark-release-recapture" cohort. To identify any impact of predictors of interest on survival in these cohorts, Cox proportional hazards models were created using the R package *coxme* v2.2-18 [15]. Interspecific models were found during diagnostics to have violated the proportional hazards assumption, and so instead separate models for each species were created within each cohort. Cox proportional hazards models including predictors of interest were compared with intercept-only "null" models by performing an analysis of variance (ANOVA) to see if inclusion significantly improved the model fit. For the multi-species cognitive experiment cohort, sex was the only candidate predictor; for the pollen-manipulation cohort, candidate predictors included eclosion mass, diet, sex, and the two-way diet:sex interaction.

##### *1.4.2 Parametric survival analyses*

The R package *flexsurv* v2.2.2 [16] was used to create parametric survival models were also created for all cohorts with the exception of the semi-natural "mark-release-recapture" cohort. These models may be described by a range of mathematical functions depending on the shape of mortality of the population, although general support for each function is mixed [17]. Intercept-only models were created by fitting survival data for each species to exponential, Weibull (Accelerated Failure Time [AFT]), Gompertz, gamma, lognormal, log-logistic, and generalised gamma distributions. For each species, the hazard functions from these (parametric) models were plotted alongside a smoothed estimate of the hazard function generated non-parametrically using the kernel density estimator from the R package *muhaz* v1.2.6.4 [18] (Supplementary Note 2). The models could then be compared visually to assess how well they fit the data, as well as statistically using their Akaike information criterion (AIC) values. The Gompertz distribution emerged as the best-supported (see Supplementary Note 2), and therefore further statistical analysis for all cohorts was carried out using parametric survival models fit to a Gompertz distribution. Cox proportional hazard models also showed no evidence for effects of sex or eclosion mass on survival, and so neither of these predictors were included in further parametric or Bayesian survival trajectory analysis.

The Gompertz function models how mortality risk changes with age, and is described by the equation $\mu(x) = \alpha e^{\beta x}$, where: $\mu(x)$ is the instantaneous mortality rate at age $x$, or the hazard, $\alpha$ is the baseline mortality independent of age, and $\beta$ is the age-dependent mortality rate (i.e. the relative change in mortality with age $x$), also known as the rate of ageing [12, 19]. A value of $\beta > 0$ reflects an increase in mortality risk with age, confirming the presence of actuarial senescence, and reflecting the process of ageing. Graphically speaking, when age is plotted against the natural log (ln) of the mortality rate (hazard), the intercept is equal to $\ln\alpha$ and the slope is equal to $\beta$, facilitating easy interpretation of these parameters: a higher intercept reflects an increase in baseline mortality risk, and a higher slope reflects an increased rate of ageing. The corresponding Gompertz survival function is given as $S(x) = e^{(\frac{\alpha}{\beta}(1-e^{\beta x})}$, where $S(x)$ is the probability of surviving to age $x$ [20]; this transformation facilitates the conversion in Figs. 1 and 2 from the parametric survival curves overlaying the empirical survival data to their corresponding log(Hazard) curves.

For the multi-species cognitive experiment cohort, 4 models were fit to the data: an intercept-only "null" model, a model including species as a predictor of just the rate parameter (representing $\alpha$, the baseline mortality), a model including species as a predictor of just the shape parameter (representing $\beta$, the rate of ageing), and a model including species as a predictor of both the rate and shape parameters. The model with the lowest AIC was selected as the best model and used for interpretation. The normboot.flexsurvreg function was used to perform bootstrapping with 1,000 iterations to generate estimates for $\alpha$ and $\beta$ for each species, and the mean and 95% confidence intervals were then extracted from these distributions. Estimates for these parameters were considered to be significantly different between species if their 95% confidence intervals did not overlap. The *H. melpomene* dataset from this cohort was subset to just those individuals surviving over 1 week; see Supplementary Note 8.

For each species in the pollen-manipulation experiment cohort, 4 models were fit to the data: an intercept-only "null" model, a model including diet as a predictor of just the rate parameter (representing $\alpha$, the baseline hazard), a model including diet as a predictor of just the shape parameter (representing $\beta$, the rate of ageing), and a model including diet as a predictor of both the rate and shape parameters. The model with the lowest AIC was selected as the best model and used for interpretation, allowing for inferences about which parameters (if any) were affected by diet treatment in either species (Supplementary Note 6). To confirm differential impact of diet on Gompertz parameters between the two species, inter-specific parametric survival models were also created which allowed either the rate parameter, the shape parameter, both, or neither, to vary by species, diet, or their interaction. The model with the lowest AIC was selected as the best model and used for interpretation, with estimates for $\alpha$ and $\beta$ parameters generated from the inter-specific model as described for the multi-species cognitive experiment cohort (Supplementary Note 6).

*1.4.3 Bayesian survival trajectory analysis*

Longevity data from the semi-natural "mark-release-recapture" cohort was analysed using the R package *BaSTA* v1.9.5 [21], which facilitates the use of incomplete recapture data to provide estimates of age-specific survival under a Bayesian framework. Though precise birth dates were known for all individuals, and the duration of the study encompassed the full lifespan of most individuals; almost no death dates were known (near-total right-censoring), and recapture probability was very low, due to the minimal-intervention approach. *BaSTA* can cope with these problems and provide estimates of important ageing parameters even with such incomplete datasets by adjusting survival estimates to recapture probabilities and using the population mean to estimate missing death dates [21, 22]. Due to the opportunistic nature of this study, data for many species suffered from low sample sizes and low recapture probabilities, which lead to wider credible intervals in *BaSTA* estimates and an underprediction of lifespan, respectively [21]. Given this, and the more thorough survival data from the other cohorts, the estimates for median lifespan for species in this cohort are almost certainly underestimates. However, these biases should apply uniformly to all species in the study. The broad patterns fit with expectations from the literature (Table 1), and we believe are still worthy of interpretation.

Birth, death, and re-sighting dates were originally recorded in days, but these were converted to 7-day bins, as cage checks were conducted on an approximately weekly basis. This allowed for a higher estimate of recapture probability, meaning models were less likely to underestimate survival [21], and led to better-fitting models with lower deviance information criterion (DIC) values (for example, dropping from 2880.91 to 2736.52 in the largest dataset, *H. melpomene*). Therefore all estimates for median lifespan and life expectancy were measured in weeks, but then manually converted back to days for ease of comparison with the other datasets presented in this chapter. This resulted in a loss of some precision in these estimates, but only in the order of +/– 1 week, and this was uniform for all species in this experiment. Similarly, estimates for Gompertz parameters were originally calculated using weeks as the unit of time, and so estimates for $\beta$ were divided by 7 to account for this in Tables 3 and S7. There were several weeks over the course of the 9-month study in which cage checks were not possible, and the reduced recapture effort for these weeks was accounted for in the models with the use of the recaptTrans argument within the basta() function.

Based on results from survival analysis of the two other cohorts, models for each species were run in *BaSTA* fit to a simple Gompertz distribution, without including sex as a covariate. Four parallel *BaSTA* simulations were run for each species, with 11,000 iterations, a burn-in of 1001, and a thinning rate of 200, to minimise serial auto-correlation. Median lifespan for each species was taken from estimates of survival probability in the *BaSTA* survQuant object. This produces estimates of survival probability for specified ages, increasing stepwise by 0.1 week each time, and so rarely included an age at which survival probability was exactly 50% (the median). Therefore median survival was taken as the mean of the two ages for which predicted survival traversed 50%. *BaSTA* also generates estimates for life expectancy as well as the Gompertz parameters b0 (equivalent to $\ln\alpha$, or ln[baseline mortality]) and b1 (equivalent to $\beta$, or age-dependent mortality rate), and these were also recorded for each species and converted to $\alpha$ and $\beta$ for ease of comparison with other cohorts. The value for maximum longevity for each species was taken as the highest age at which an individual of that species was observed alive (Table 3). Correlation between median lifespan estimates from *BaSTA* and existing reported maximum lifespans (Table 1) was tested using Pearson's correlation test; see Supplementary Note 5 for a plot of this correlation.

*Heliconius himera*, *Philaethria dido*, and *Heliconius charithonia* were excluded from *BaSTA*s due to low sample size and few re-sightings, and so maximum longevity was the only metric retained for these species. *BaSTA* also does not allow for recaptures during the "birth week", and so 66 sightings which occurred in the first week of an individual's life were excluded from analysis. Accounting for these exclusions, *BaSTA*s were performed using data from a total of 941 butterflies, with 959 recorded sightings of 378 different individuals. A full breakdown of these exclusions per species can be found in Supplementary Note 5.

##### *1.4.4 Feeding habit differences*

The maximum reported lifespans from Table 1 as well as the final parameters generated from the multi-species cognitive experiment cohort and the semi-natural "mark-release-recapture" cohort were then analysed for broad differences between pollen-feeders

(*Heliconius* species) versus non-pollen-feeders (the outgroups). This was first done using standard statistical techniques, treating each species as an independent observation.

The Shapiro-Wilk test of normality showed all parameters derived from the multi-species cognitive experiment cohort to be normally distributed (median lifespan: $W = 0.92$, $p = 0.530$; maximum lifespan: $W = 0.89$, $p = 0.382$; Gompertz parameter $\alpha$: $W = 0.83$, $p = 0.129$; Gompertz parameter $\beta$: $W = 0.97$, $p = 0.862$), and the F-test of equality of variances showed all measures to have equal variances between the two groups (median lifespan: $F_{2,1} = 1.95$, $p = 0.903$; maximum lifespan: $F_{2,1} = 93.17$, $p = 0.146$; $\alpha$: $F_{2,1} = 86.95$, $p = 0.151$; $\beta$: $F_{2,1} = 3.75$, $p = 0.686$), and so these were assessed using a two-tailed Student's t-test to test for differences between feeding habits.

The Shapiro-Wilk test of normality showed median and maximum lifespans derived from the semi-natural "mark-release-recapture" cohort to be normally distributed (median: $W = 0.94$, $p = 0.335$; maximum: $W = 0.93$, $p = 0.214$), and the F-test of equality of variances showed both measures to have equal variances between the two groups (median: $F_{11,4} = 5.24$, $p = 0.124$; maximum: $F_{11,4} = 4.90$, $p = 0.214$), and so these were assessed using a two-tailed Student's t-test to test for differences between feeding habits. The Shapiro-Wilk test showed the Gompertz parameters $\alpha$ and $\beta$ derived from results in this cohort to be non-normally distributed ($\alpha$: $W = 0.59$, $p < 0.001$; $\beta$: $W = 0.73$, $p < 0.001$), so these were assessed using a two-tailed Mann-Whitney U test to test for differences between feeding habits.

The Shapiro-Wilk test showed maximum lifespan values from Table 1 to be normally distributed ($W = 0.96$, $p = 0.298$), and the F-test of equality of variances this measure to have unequal variances between the two groups ($F_{20,5} = 6.72$, $p = 0.044$) and so these were assessed using a two-tailed Welch's t-test to test for differences between feeding habits.

Although pollen feeding is very tightly confounded by phylogenetic relatedness in our dataset, we explored the potential to remove phylogenetic effects by repeating these comparisons using the phylANOVA() function from the R package *phytools* v2.3-0 [23], run with 10,000 simulations using a trimmed phylogenetic tree taken from [24]. This package and tree were also used to create the phylogenetic tree in Fig. 1, and to estimate phylogenetic signal as measured by Pagel's λ. Results from these phylogenetic ANOVAs are presented in Supplementary Note 3 for completeness.

*1.4.5 Functional senescence analyses*

For all functional senescence analyses from the pollen-manipulation experiment cohort, general or generalised linear mixed models were fit, for two single-species datasets containing data for each species across their full lifespans (up to week 5 for *D. iulia* and up to week 17 for *H. hecale*), as well as for an interspecific dataset containing data for *H. hecale* and *D. iulia* up to week 5 (the oldest age at which there was data for *D. iulia*). All models included individual ID as a random effect due to the repeated-measures nature of the analysis. All single-species "full" models included diet, age, sex, and their three-way interaction as candidate predictors. All interspecific "full" models included species, diet, age, and their three-way interaction as candidate predictors, as well as two-way interactions between species and any candidate predictors found to be significant in the single-species models. All models also included longevity as a candidate predictor, to account for the possibility of selective disappearance and ensure unbiased estimates of the effect of age. Other candidate predictors tested in the initial "full" model depended on the assay, and were selected based on biological relevance; these are detailed below.

For analysis of body mass and grip strength data, linear mixed models were fit using the R package *lme4* v1.1-34 [25] using Satterthwaite's approximation from the package *lmerTest* 3.1-3 [26]. For body mass, additional candidate predictors included assay time (measured as the fraction of the day elapsed since midnight). For grip strength, additional candidate predictors included assay time and eclosion mass (measured in grams [g]). For analysis of flight behaviour data, generalised linear mixed models were fit with an *ordbeta* distribution [27], based on recommendations for the analysis of continuous proportion data [28], using the R package *glmmTMB* v1.1.7 [29]. Additional candidate predictors included forewing length (measured in millimetres [mm]), illuminance (measured in lux), assay time, whether it was raining, and whether the butterfly had an intact wing apex (used as an index of wing-wear). Values for illuminance were centred and scaled to aid model convergence.

For all models, stepwise backward model selection was implemented by performing an ANOVA on the model with and without a named predictor, to see if inclusion of the predictor significantly improved model fit. The final "best" model was then compared with an intercept-only "null" model to confirm its validity, also using an ANOVA. Age, diet, and species (for the inter-specific models) were always included in the final model as these were effects of interest, as was longevity, which ensured unbiased age estimates. An ANOVA was then run on the final model to report significance of named predictors. If an effect was not included in the final model, the nonsignificant result of the ANOVA from model selection resulting in its removal was instead reported. Diagnostics for all models were performed using the R package *DHARMa* v0.4.6 [30]. Model predictions for visualisation of results were created with the aid of the R packages *effects* v4.2-2 [31] and *sjPlot* v2.8.15 [32].

#### 2. Supplementary Notes

##### *2.1 Supplementary Note 1: Additional results on flight behaviour in ageing* **H. hecale** *and* **D. iulia**

While *H. hecale* spent less time flying later in the day ($\chi^2_1$ = 3.95, *p* = 0.047), in contrast to the grip strength assay, neither age ($\chi^2_1$ = 2.50, *p* = 0.114) nor diet ($\chi^2_1$ = 0.11, *p* = 0.735) was found to be a significant predictor of time spent in active flight in this species (Fig. S3). In *D. iulia*, there was an interaction between age, sex, and diet ($\chi^2_1$ = 4.89, *p* = 0.027) such that in females, pollen-fed butterflies spent more time in active flight as they aged, whereas pollen-deprived butterflies spent less time in active flight as they aged (Fig. S3); an unexpected result perhaps indicative of a sex-specific response to the presence of natural floral cues, which would need further investigation to interrogate fully. Environment also had an impact on flight behaviour in *D. iulia*, with butterflies spending less time flying if it was raining ($\chi^2_1$ = 5.18, *p* = 0.023). Broadly, these results suggest a lack of senescence in either butterfly in the flight activity assay, which is surprising considering the evidence for reduced flight capacity with age in other insects [5], including *Drosophila* [4] and the butterfly *Pieris napi* [6]. This is likely because in the short (45-second) assay presented here, differences in flight activity seem to reflect behavioural choices rather than physiological capacity, given that individuals flew less later in the day (*H. hecale*), or when it was raining (*D. iulia*) – behaviour which aligns with common knowledge about when these butterflies are most active [33]. Considering that many individuals were spotted flying vigorously around the cages just hours before being found dead (personal observation), it seems unlikely that a reduction in total flight capacity in ageing individuals would be observed within the 45-second assay. These results do not preclude the possibility that observed reductions in flight at the metabolic level in other butterflies [34, 35] might be also recapitulated in Heliconiini. Further investigation would help to determine if similar patterns of differential senescence between *Heliconius* and the shorter-lived Heliconiini may be identified in other aspects of their physiology.

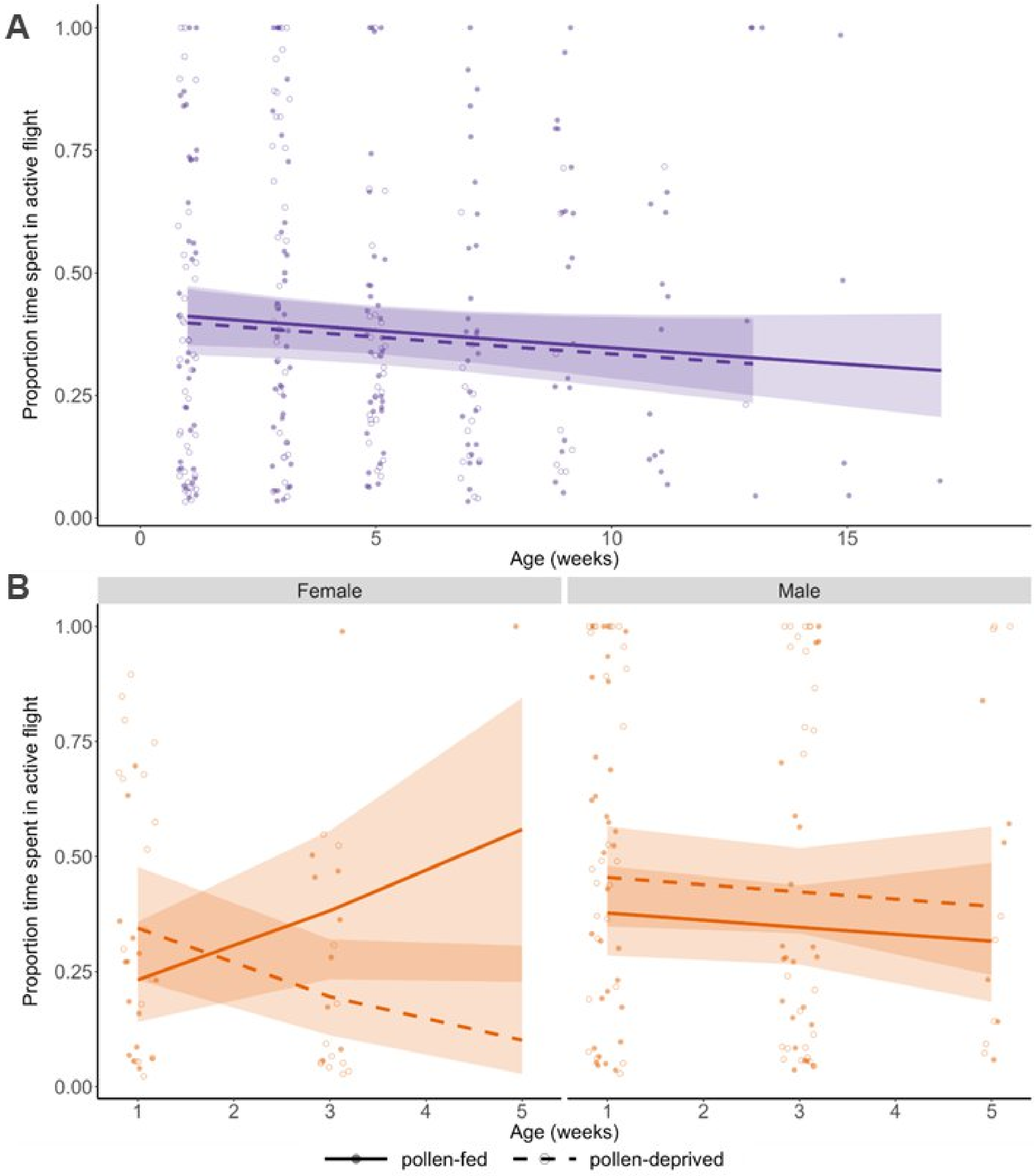

**Fig. S3: Flight behaviour results.**

Dots represent individual data points indicating the proportion of time an individual spent in active flight during a 45-second assay period, for both for (A) *H. hecale* and (B) *D. iulia.* Regression lines, with 95% confidence intervals, are from GLMM analysis and show predicted proportion of time spent in active flight for each species / diet combination, with the sexes separated into facets for *D. iulia* due to the presence of a three-way age:sex:diet interaction.

##### *2.2 Supplementary Note 2: Hazard function distribution selection*

Support for distributions was mixed, and the best-fitting distribution both visually and in terms of the lowest AIC value was different for different species (Figs. S4 and S5; Tables S1 and S2). This may be in part due to sample size, as some sources suggest a minimum of 100 individuals per group to distinguish between mortality functions [20], which was not true for all groups analysed, though groups did largely reach the suggested minimum of 50 individuals each to estimate age-specific mortality without introducing significant bias [36]. Choosing a distribution based on prior knowledge was also problematic, with few studies focusing on actuarial senescence in Lepidoptera; but those that do indicate that the Gompertz and Weibull models are among the best-fitting [37-39]. Considering all available information, the Gompertz distribution emerged as the best-supported overall. In particular, data from the pollen-manipulation experiment cohort, which is likely the best reflection of natural senescence, fit the Gompertz distribution extremely well (Fig. S5, Table S2). Survival data from the multi-species cognitive experiment cohort, while useful, is likely to more poorly reflect natural ageing patterns considering the strenuous conditions of this experiment, where survival was conditional upon learning a colour association, and butterflies were subjected to frequent handling and regular bouts of food deprivation.

**Table S1:** Comparison of Akaike information criterion (AIC) values for parametric survival models fit to various mortality distributions for survival data from the multi-species cognitive experiment cohort. Distributions for each species are presented in order of most- to least-supported. Weibull (AFT) = Weibull (accelerated failure time).

| Species | Distribution | ΔAIC | Degrees of freedom |
|---|---|---|---|
| *A. vanillae* | Weibull (AFT) | 0 | 2 |
| | Gompertz | 0.62 | 2 |
| | Gamma | 1.01 | 2 |
| | Generalised gamma | 1.35 | 3 |
| | Lognormal | 9.65 | 2 |
| | Log-logistic | 20.41 | 2 |
| | Exponential | 26.87 | 1 |
| *D. phaetusa* | Generalised gamma | 0 | 3 |
| | Gompertz | 0.05 | 2 |
| | Weibull (AFT) | 21.89 | 2 |
| | Gamma | 35.03 | 2 |
| | Log-logistic | 55.07 | 2 |
| | Lognormal | 58.30 | 2 |
| | Exponential | 67.77 | 1 |
| *D. iulia* | Weibull (AFT) | 0 | 2 |
| | Generalised gamma | 0.57 | 3 |
| | Gamma | 5.40 | 2 |
| | Gompertz | 7.70 | 2 |
| | Lognormal | 32.36 | 2 |
| | Log-logistic | 33.65 | 2 |
| | Exponential | 60.19 | 1 |
| *H. hecale* | Generalised gamma | 0 | 3 |
| | Gompertz | 8.12 | 2 |
| | Exponential | 18.53 | 1 |
| | Weibull (AFT) | 19.61 | 2 |
| | Gamma | 20.35 | 2 |
| | Log-logistic | 32.34 | 2 |
| | Lognormal | 33.57 | 2 |
| *H. melpomene* | Weibull (AFT) | 0 | 2 |
| | Gamma | 0.61 | 2 |
| | Log-logistic | 1.62 | 2 |
| | Generalised gamma | 1.98 | 3 |
| | Gompertz | 2.44 | 2 |
| | Lognormal | 3.53 | 2 |
| | Exponential | 15.01 | 1 |

**Table S2:** Comparison of Akaike information criterion (AIC) values for parametric survival models fit to various mortality distributions for survival data from the pollen-manipulation experiment cohort. Distributions for each species are presented in order of most- to least-supported. Weibull (AFT) = Weibull (accelerated failure time).

| Species | Distribution | ΔAIC | Degrees of freedom |
|---|---|---|---|
| *H. hecale* | Gompertz | 0 | 2 |
| | Generalised gamma | 3.51 | 3 |
| | Weibull (AFT) | 9.87 | 2 |
| | Gamma | 15.97 | 2 |
| | Exponential | 24.01 | 1 |
| | Log-logistic | 28.46 | 2 |
| | Lognormal | 42.53 | 2 |
| *D. iulia* | Gompertz | 0 | 2 |
| | Generalised gamma | 4.04 | 3 |
| | Weibull (AFT) | 11.21 | 2 |
| | Gamma | 24.09 | 2 |
| | Log-logistic | 31.86 | 2 |
| | Lognormal | 49.12 | 2 |
| | Exponential | 62.77 | 1 |

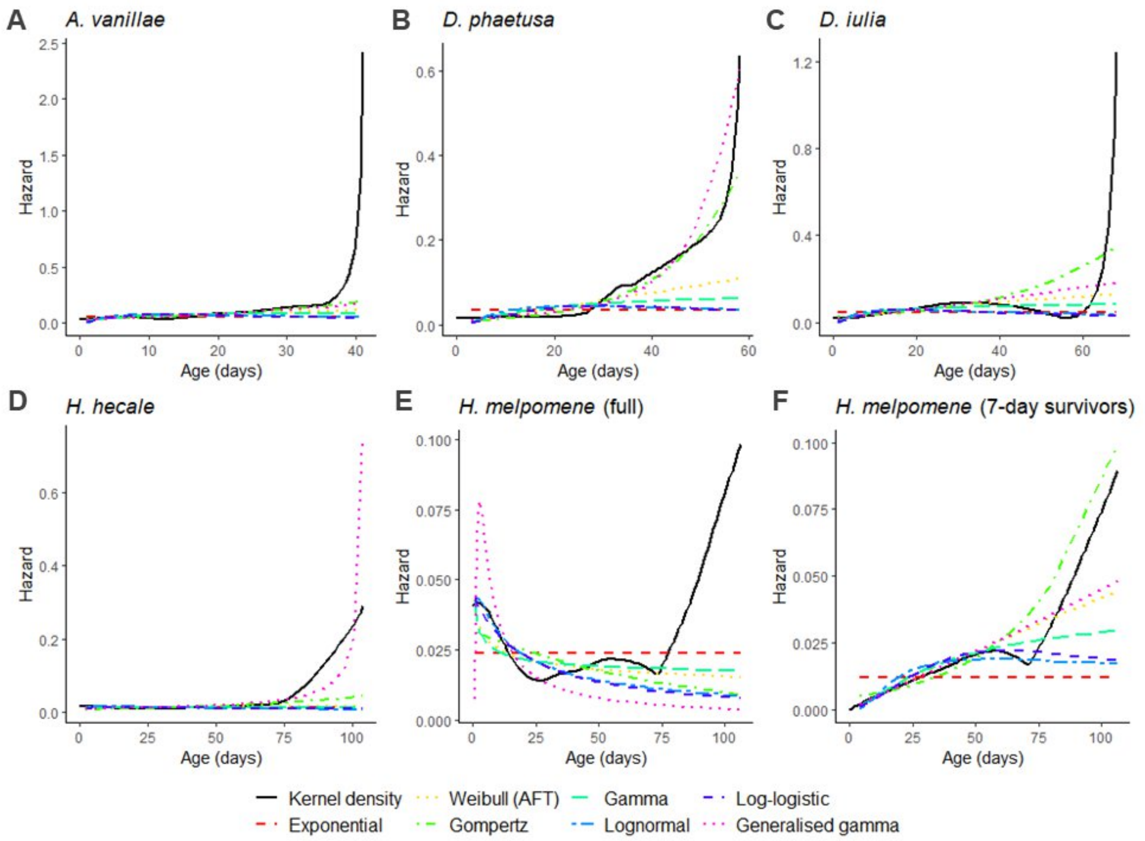

**Fig. S4: Hazard rate curves fit to various mortality distributions for the multi-species cognitive experiment cohort.**

The kernel density estimate is used to compare fit of the various models to the actual data from the experiment. Both the full *H. melpomene* dataset (E) and the 7-day survivors subset (F) are plotted to justify the use of the latter for further statistical analysis (see Supplementary Note 7).

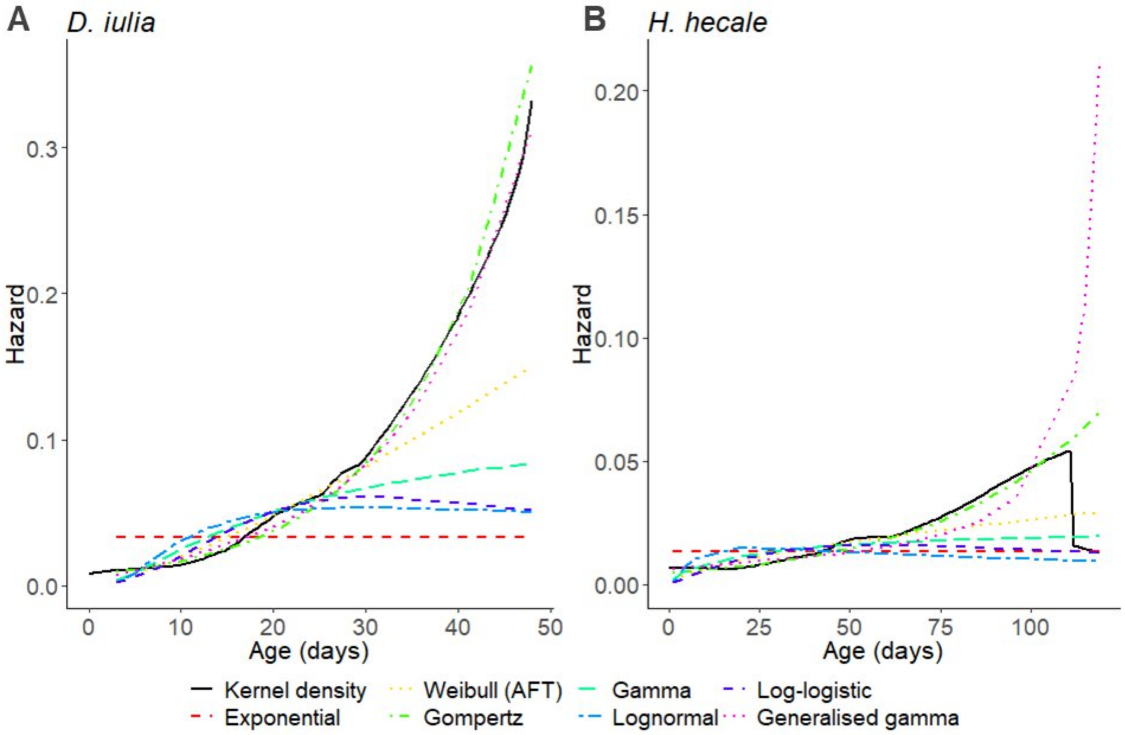

**Fig. S5: Hazard rate curves fit to various mortality distributions for butterflies in the pollen-manipulation experiment cohort.**

The kernel density estimate is used to compare fit of the various models to the actual data from the experiment.

##### *2.3 Supplementary Note 3: Results of comparisons between feeding habits when controlling for phylogenetic relatedness*

In order to control for phylogenetic relatedness, all comparisons of ageing parameters between feeding habit (pollen-feeding versus non-pollen-feeding) were then tested using a phylogenetic ANOVA. This rendered the difference between feeding habits in maximum reported lifespans (Table 1) non-significant ($F = 13.18$, $p = 0.212$), while the difference between feeding habits in median ($F = 19.16$, $p = 0.078$) and maximum ($F = 23.39$, $p = 0.064$) lifespans from the smaller multi-species cognitive experiment cohort also showed non-significant trends. When applied to the larger semi-natural "mark-release-recapture" cohort, the difference between feeding habits in maximum lifespan was no longer significant ($F = 9.31$, $p = 0.240$). We conclude that because this evolutionary transition to pollen-feeding only occurred once at the base of a monophyletic clade, there is insufficient information to disentangle effects of phylogeny and pollen-feeding. It is notable, however, that differences between feeding habits in median lifespan estimates ($F = 36.28$, $p = 0.03$) and baseline mortality ($\alpha$) ($F = 29.57$, $p = 0.048$) in the semi-natural "mark-release-recapture" cohort remained significant even when controlling for phylogenetic relatedness, while the trends in other analyses are consistent with our wider interpretations. As such, while we cannot discount phylogenetic effects as a factor in our analyses, the pronounced shifts we observed are unlikely to have evolved under neutral evolution.

##### *2.4 Supplementary Note 4: Details of parametric survival analysis for the multi-species cognitive experiment cohort*

The best-supported parametric survival model for the multi-species cognitive experiment cohort allowed both baseline mortality ($\alpha$) and the rate of ageing ($\beta$) to vary by species (Table S3). Both parameters showed trends towards reduced ageing in pollen-feeding *Heliconius*, with baseline mortality ($\alpha$) highest in *A. vanillae* (non-pollen-feeding), followed by *D. iulia* (non-pollen-feeding), *D. phaetusa* (non-pollen-feeding), *H. hecale* (pollen-feeding), and finally *H. melpomene* (pollen-feeding); however, statistical differences did not partition with respect to feeding habit, with baseline mortality ($\alpha$) for *A. vanillae* and *D. iulia* significantly higher to that of *D. phaetusa*, *H. hecale*, and *H. melpomene* (see Table S4 for details of overlapping confidence intervals and Table S5 for full contrasts). Similarly, rate of ageing ($\beta$) was highest in *D. phaetusa* (non-pollen-feeding)¸ followed by *A. vanillae* (non-pollen-feeding), *D. iulia* (non-pollen-feeding), *H. melpomene* (pollen-feeding), and finally *H. hecale* (pollen-feeding); however, the only statistically significant differences were between *D. phaetusa* versus *D. iulia*, *H. hecale*, and *H. melpomene*; as well as between *H. hecale* versus both *A. vanillae* and *D. iulia* (Tables S4 and S6). When parameters were pooled for pollen-feeders ($n$ = 2) and non-pollen-feeders ($n$ = 3), neither baseline mortality ($\alpha$) ($t_3$ = 1.60, $p$ = 0.208) nor the rate of ageing ($\beta$) ($t_3$ = 2.30, $p$ = 0.105) segregated statistically by feeding habit, likely related to the lack of power in this comparison. Despite this, both median and maximum lifespan segregated by feeding habit in statistical analysis (see Results).

**Table S3:** Comparison with Akaike information criterion (AIC) of parametric survival models fit to data from each species in the multi-species cognitive experiment cohort using a Gompertz distribution. Different models allowed either the rate parameter, the shape parameter, both, or neither, to vary by species. The rate parameter is the Gompertz parameter α, or baseline mortality, and the shape parameter is the Gompertz parameter β, or rate of ageing.

| Model specification | ΔAIC | Degrees of freedom |
|---|---|---|
| rate~Species, shape~Species | 0 | 10 |
| rate~Species, shape~1 | 24.12 | 6 |
| rate~1, shape~Species | 68.63 | 6 |
| rate~1, shape~1 | 344.39 | 2 |

**Table S4:** Bootstrapped parameter estimates and confidence intervals for each species in the multispecies cognitive experiment cohort, based on the final model: rate~Species, shape~Species. Results here are presented to 2 significant figures rather than 2 decimal places due to the small size of several estimates. CI = confidence interval.

| **Species** | **Feeding habit** | **Parameter** | **Parameter estimate** | **Lower 95% CI** | **Upper 95% CI** |
|---|---|---|---|---|---|
| *A. vanillae* | non-pollen-feeding | α – baseline mortality | 0.032888 | 0.024787 | 0.043253 |
| | | β – rate of ageing | 0.043864 | 0.027995 | 0.058818 |
| *D. phaetusa* | non-pollen-feeding | α – baseline mortality | 0.00746 | 0.004405 | 0.011608 |
| | | β – rate of ageing | 0.067497 | 0.05166 | 0.083497 |
| *D. iulia* | non-pollen-feeding | α – baseline mortality | 0.023793 | 0.018824 | 0.029448 |
| | | β – rate of ageing | 0.039938 | 0.030494 | 0.049916 |
| *H. hecale* | pollen-feeding | α – baseline mortality | 0.006959 | 0.004133 | 0.010645 |
| | | β – rate of ageing | 0.017819 | 0.008539 | 0.027959 |
| *H. melpomene* | pollen-feeding | α – baseline mortality | 0.005005 | 0.002372 | 0.009322 |
| | | β – rate of ageing | 0.028706 | 0.014067 | 0.042447 |

**Table S5:** Inter-species contrasts for estimates of the α / rate parameter (baseline mortality) in the final inter-specific model for the multi-species cognitive experiment cohort: rate~Species, shape~Species. An asterisk denotes significant differences between pairs of species, based on non-overlapping confidence intervals as shown in Table S4. PF = pollen-feeding, NPF = non-pollen-feeding.

| | *A. vanillae* (NPF) | *D. phaetusa* (NPF) | *D. iulia* (NPF) | *H. hecale* (PF) | *H. melpomene* (PF) |
|---|---|---|---|---|---|
| *A. vanillae* (NPF) | | * | | * | * |
| *D. phaetusa* (NPF) | | | * | | |
| *D. iulia* (NPF) | | | | * | * |
| *H. hecale* (PF) | | | | | |
| *H. melpomene* (PF) | | | | | |

**Table S6:** Inter-species contrasts for estimates of the β / shape parameter (rate of ageing) in the final inter-specific model for the multi-species cognitive experiment cohort: rate~Species, shape~Species. An asterisk denotes significant differences between pairs of species, based on non-overlapping confidence intervals as shown in Table S4. PF = pollen-feeding, NPF = non-pollen-feeding.

| | *A. vanillae* (NPF) | *D. phaetusa* (NPF) | *D. iulia* (NPF) | *H. hecale* (PF) | *H. melpomene* (PF) |
|---|---|---|---|---|---|
| *A. vanillae* (NPF) | | | | * | |
| *D. phaetusa* (NPF) | | | * | * | * |
| *D. iulia* (NPF) | | | | * | |
| *H. hecale* (PF) | | | | | |
| *H. melpomene* (PF) | | | | | |

##### *2.5 Supplementary Note 5: Details of survival analysis for the semi-natural "mark-release-recapture" cohort*

Results from Bayesian survival trajectory analysis on the semi-natural "mark-release-recapture" cohort showed lower baseline mortality ($\alpha$) in pollen-feeding species ($W = 0$, $p < 0.001$), but not rate of ageing ($\beta$) ($W = 17$, $p = 0.195$), although this trended lower in non-pollen-feeders. These results are presented graphically here to aid interpretation (Fig. S6). Estimates for median lifespan from this cohort are also presented graphically here (Fig. S6) and appear to be underestimated when compared with survival data from the other, more closely-tracked cohorts (Tables 2 and 4). However, these showed a strong positive correlation with maximum reported lifespan for each species as presented in Table 1 ($r = 0.64$; $t_{15} = 3.25$, $p = 0.005$). The correlation plot is also presented here to aid interpretation (Fig. S6). Full details of sample size and parameter estimates for this cohort are presented in Table S7.

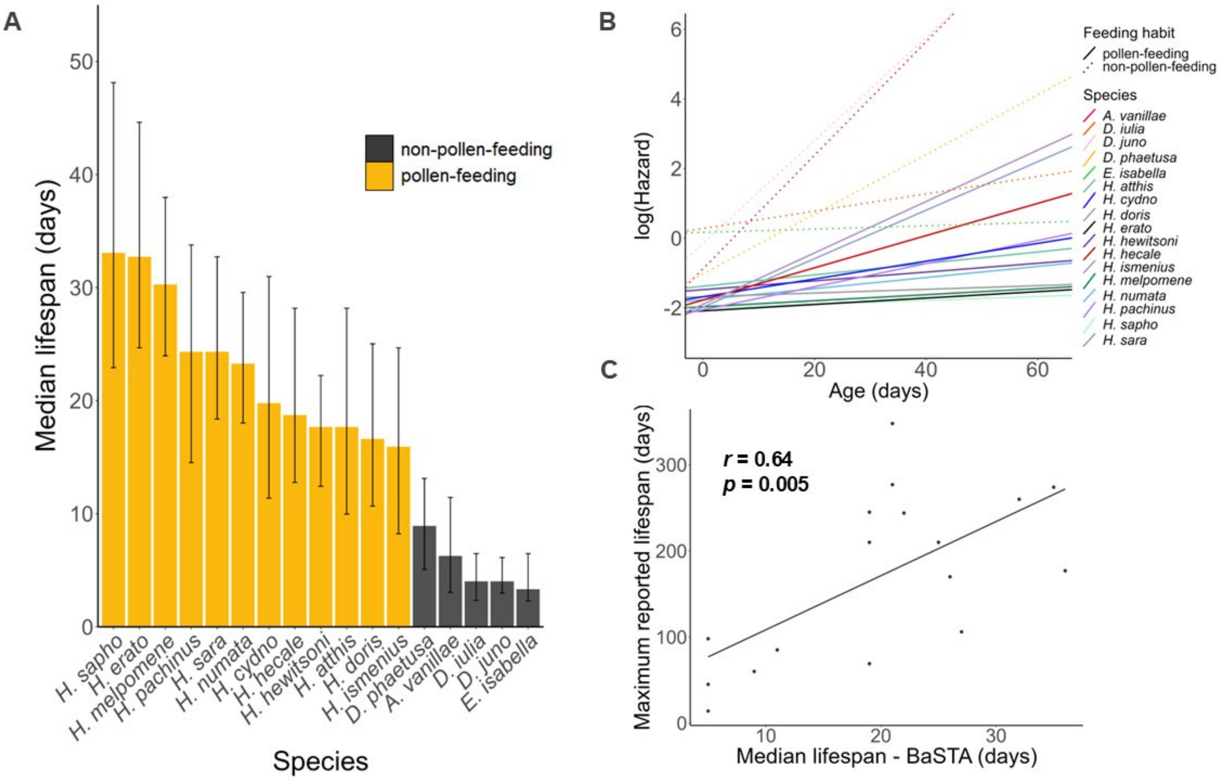

**Fig. S6: Results of *BaSTA* analysis in the semi-natural "mark-release-recapture" cohort and their relationship to existing lifespan data.**

**A** Median lifespan estimates with 95% predictive intervals for each species in the cohort, coloured according to feeding habit. **B** log(Hazard) curves fit to a Gompertz distribution for all species in the semi-natural "mark-release-recapture" cohort, based on estimates for α and β derived from *BaSTA* analysis (see Table S7 for full details). An increase in the intercept on this graph represents an increase in baseline mortality (α); an increase in the slope represents an increase in the rate of ageing (β). **C** Correlation between median lifespan estimates derived from *BaSTA* analysis for species in this cohort and maximum reported lifespans for each species as listed in Table 1. *r* = Pearson's correlation coefficient.

**Table S7: Sample size, recorded sightings, and longevity metrics from display cage *BaSTA* results.**

Species are presented in order of descending *n* sighted (original). *N* placed refers to the number of individuals marked and placed into the display cage for the study. *n* sighted (original) refers to the number of individuals that were re-sighted at least once, and *n* sighted (retained) refers to the number of these retained after excluding those sighted within the first week of life (see Supplementary Methods for details). Total detections refers to the total number of sightings for that species (many individuals were re-sighted more than once). Non-pollen-feeders are coloured in red for emphasis (see Table 3). *H. himera*, *P. dido*, and *H. charithonia* were excluded from the analysis due to low sample size. Results for β are presented to 2 significant figures rather than 2 decimal places due to the small size of several estimates. CI = credible interval, PI = predictive interval.

| Species | *N* placed | *n* sighted (original) | *n* sighted (retained) | Total detections | α | | | β | | | Median lifespan (days) | | | Maximum recorded lifespan (days) |
|---|---|---|---|---|---|---|---|---|---|---|---|---|---|---|
| | | | | | estimate | lower 95% CI | upper 95% CI | estimate | lower 95% CI | upper 95% CI | estimate | lower 95% PI | upper 95% PI | |
| *H. melpomene* | 285 | 116 | 90 | 261 | 0.14 | 0.10 | 0.19 | 0.0087 | 0.0018 | 0.016 | 30.28 | 23.98 | 37.98 | 102 |
| *H. numata* | 94 | 65 | 63 | 110 | 0.17 | 0.12 | 0.25 | 0.016 | 0.0053 | 0.025 | 23.28 | 18.03 | 29.58 | 106 |
| *H. sapho* | 81 | 50 | 40 | 183 | 0.13 | 0.09 | 0.19 | 0.0056 | 0.00038 | 0.012 | 32.73 | 24.68 | 44.63 | 143 |
| *H. sara* | 70 | 49 | 45 | 134 | 0.19 | 0.13 | 0.26 | 0.0053 | 0.00045 | 0.013 | 24.33 | 18.38 | 32.73 | 107 |
| *H. erato* | 46 | 25 | 21 | 43 | 0.12 | 0.06 | 0.21 | 0.0092 | 0.00061 | 0.023 | 33.08 | 22.93 | 48.13 | 94 |
| *H. doris* | 32 | 24 | 22 | 51 | 0.14 | 0.05 | 0.31 | 0.070 | 0.022 | 0.12 | 17.68 | 12.43 | 22.23 | 35 |
| *H. hecale* | 47 | 21 | 18 | 33 | 0.23 | 0.12 | 0.37 | 0.013 | 0.00077 | 0.032 | 18.73 | 12.78 | 28.18 | 78 |
| *H. atthis* | 33 | 16 | 15 | 34 | 0.25 | 0.11 | 0.45 | 0.016 | 0.00069 | 0.043 | 16.63 | 10.68 | 25.03 | 53 |
| *H. pachinus* | 19 | 14 | 14 | 31 | 0.13 | 0.05 | 0.31 | 0.033 | 0.0065 | 0.066 | 24.33 | 14.53 | 33.78 | 58 |
| *H. cydno* | 19 | 13 | 9 | 23 | 0.19 | 0.06 | 0.41 | 0.026 | 0.0021 | 0.060 | 19.78 | 11.38 | 30.98 | 55 |
| *D. phaetusa* | 30 | 12 | 10 | 14 | 0.36 | 0.10 | 0.91 | 0.086 | 0.010 | 0.19 | 8.93 | 5.08 | 13.13 | 17 |
| *D. juno* | 45 | 9 | 5 | 5 | 0.86 | 0.22 | 1.91 | 0.15 | 0.0097 | 0.38 | 4.03 | 2.35 | 6.48 | 11 |
| *H. hewitsoni* | 15 | 9 | 9 | 13 | 0.17 | 0.03 | 0.46 | 0.046 | 0.0051 | 0.12 | 17.68 | 9.98 | 28.18 | 41 |
| *E. isabella* | 59 | 8 | 4 | 4 | 1.18 | 0.79 | 1.74 | 0.0047 | 0.00010 | 0.016 | 4.03 | 2.98 | 6.13 | 50 |
| *H. ismenius* | 20 | 8 | 8 | 14 | 0.15 | 0.01 | 0.54 | 0.074 | 0.0054 | 0.19 | 15.93 | 8.23 | 24.68 | 24 |
| *A. vanillae* | 12 | 3 | 3 | 3 | 0.42 | 0.07 | 1.51 | 0.16 | 0.012 | 0.36 | 6.27 | 3.05 | 11.45 | 11 |
| *D. iulia* | 34 | 2 | 2 | 3 | 1.32 | 0.63 | 2.29 | 0.025 | 0.00032 | 0.086 | 3.33 | 2.28 | 6.48 | 25 |
| *H. himera* | 2 | 2 | | | | | | | | | | | | 17 |
| *P. dido* | 13 | 1 | | | | | | | | | | | | 7 |
| *H. charithonia* | 3 | 1 | | | | | | | | | | | | 22 |

#### *2.6 Supplementary Note 6: Details of parametric survival analysis for the pollen-manipulation experiment cohort*

The best-supported inter-specific parametric survival model for the pollen-manipulation experiment cohort allowed the rate of ageing ($\beta$) to vary by species, and baseline mortality ($\alpha$) to vary by a two-way species:diet interaction (Table S8). The meaning of this interaction is illuminated by the two single-species models, where the best supported model in *H. hecale*, but not *D. iulia*, to vary by diet (Table S9). Table S10 shows bootstrapped parameter estimates derived from the inter-specific model for each species/diet combination.

**Table S8:** Comparison with Akaike information criterion (AIC) of parametric survival models fit to data from both species in the pollen-manipulation experiment cohort using a Gompertz distribution. Different models allowed either the rate parameter, the shape parameter, both, or neither, to vary by species, diet, and their two-way interaction. The rate parameter is the Gompertz parameter α, or baseline mortality, and the shape parameter is the Gompertz parameter β, or rate of ageing.

| Model specification | ΔAIC | Degrees of freedom |
|---|---|---|
| rate~Species*Diet, shape~Species | 0 | 6 |
| rate~Species*Diet, shape~Species+Diet | 1.52 | 7 |
| rate~Species*Diet, shape~Species*Diet | 3.50 | 8 |
| rate~Species*Diet, shape~1 | 25.14 | 5 |
| rate~Species*Diet, shape~Diet | 26.26 | 6 |
| rate~Species+Diet, shape~1 | 31.26 | 4 |
| rate~Species, shape~1 | 33.15 | 3 |
| rate~Diet, shape~1 | 83.54 | 3 |
| rate~1, shape~1 | 86.52 | 2 |

**Table S9:** Comparison with Akaike information criterion (AIC) of parametric survival models fit to data from both species in the pollen-manipulation cohort using a Gompertz distribution. Different models allowed either the rate parameter, the shape parameter, both, or neither, to vary by diet. The rate parameter is the Gompertz parameter α, or baseline mortality, and the shape parameter is the Gompertz parameter β, or rate of ageing.

| Species | Model specification | ΔAIC | Degrees of freedom |
|---|---|---|---|
| *H. hecale* | rate~Diet, shape~1 | 0 | 3 |
| | rate~1, shape~Diet | 0.32 | 3 |
| | rate~Diet, shape~Diet | 1.52 | 4 |
| | rate~1, shape~1 | 6.10 | 2 |
| *D. iulia* | rate~1, shape~1 | 0 | 2 |
| | rate~Diet, shape~1 | 1.90 | 3 |
| | rate~1, shape~Diet | 1.95 | 3 |
| | rate~Diet, shape~Diet | 3.88 | 4 |

**Table S10:** Bootstrapped parameter estimates and confidence intervals for each species/diet combination in the pollen-manipulation experiment cohort, based on the final model: rate~Species*Diet, shape~Species. Results here are presented to 2 significant figures rather than 2 decimal places due to the small size of several estimates. CI = confidence interval.

| Species | Diet | Parameter | Parameter estimate | Lower 95% CI | Upper 95% CI |
|---|---|---|---|---|---|
| *H. hecale* | pollen-fed | α | 0.0044 | 0.0023 | 0.0076 |
| | | β | 0.025 | 0.016 | 0.034 |
| | pollen-deprived | α | 0.0087 | 0.0052 | 0.014 |
| | | β | 0.025 | 0.016 | 0.034 |
| *D. iulia* | pollen-fed | α | 0.0077 | 0.0042 | 0.012 |
| | | β | 0.081 | 0.063 | 0.10 |
| | pollen-deprived | α | 0.0072 | 0.0039 | 0.012 |
| | | β | 0.081 | 0.063 | 0.10 |

##### *2.7 Supplementary Note 7: Maximum reported lifespans from butterfly exhibitors*

For many species, multiple records of maximum longevity were found, sourced from data from butterfly exhibitors, mark-release-recapture studies, and insectary populations (Table 1; Supplementary Data 1). In all such instances, maximum lifespan records from butterfly exhibitors eclipsed those from insectary populations and mark-release recapture studies in the wild. One dataset in particular (Kelson, unpublished) was responsible for many of the highest maximum lifespan records for various species. This was provided by Mr. Richard Kelson, the entomologist running the Butterfly Habitat at Six Flags Discovery Kingdom in Vallejo, California, who has been compiling a dataset of maximum lifespan for over 180 butterfly species kept at the exhibit since 1989. The methodology for this data collection involves a) ordering pupae in batches, and recording when the last individual of each species ecloses; b) conducting a weekly species census to know which species are still alive, being especially careful to note sightings for species with only a few remaining living individuals; and c) calculating maximum longevity by subtracting the latest possible adult eclosion date from the last date an individual of that species was seen alive (so this dataset is right-censored). This dataset includes many non-Heliconiini butterfly species, and an up-to-date version presented in full in Supplementary Data 2 for other researchers interested in butterfly lifespan. Species are listed alphabetically according to names used by the U.S. Department of Agriculture; where current taxonomic names are different, these are reported in parentheses. References to Watts, 2004 [40] are made where that study found greater lifespans.

#### *2.8 Supplementary Note 8:* **H. melpomene** *early deaths exclusion*

The learning and memory assays performed on individuals from the multi-species cognitive experiment cohort necessitated frequent handling of the butterflies, as well as regular bouts of food-deprivation during the colour choice assays, introducing stressors which likely impacted survival. One example of this was the high early mortality demonstrated in *H. melpomene*, with a substantial number of individuals ($n$ = 37; 35.92%) dying in the first week. Inclusion of this early spike in mortality in survival analysis greatly biased estimates of median survival time for this species, and also produced uninformative estimates of other ageing parameters, including a conclusion of negative senescence which is not supported by any other data in this paper (Fig. S7). Visual inspection confirmed that removal of these early deaths allowed a much better fit of the data to standard parametric survival distributions (Fig. S4). This incongruity fits with the hypothesis that the high early mortality observed was likely due to an inability to adapt to the contrived conditions in the experimental cages, rather than a reflection of senescence in natural conditions. Therefore only individuals of *H. melpomene* that survived for longer than a week were included in further statistical analysis ($n$ = 66, with 31 censored observations). Subsetting the other species in this dataset to just individuals who survived the first 7 days of the experiments resulted in qualitatively similar findings (Fig. 1, Fig. S7). However, there was no support for performing this exclusion in these species (Fig. S4), and so further statistical analysis for all other species was carried out on the full group.

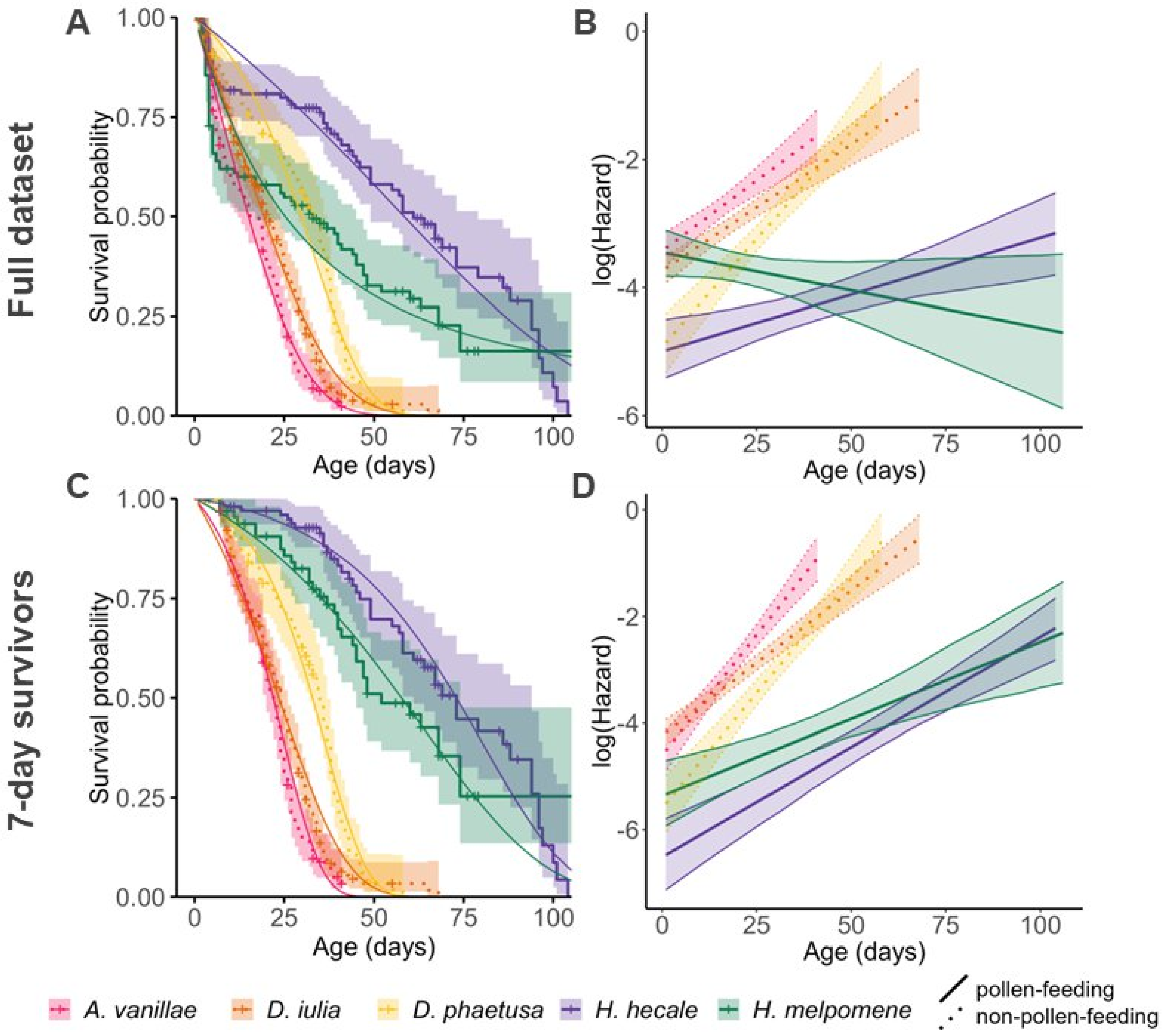

**Fig. S7: Survival and log(Hazard) curves for butterflies in the multi-species cognitive experiment cohort for (A-B) the full dataset and (C-D) only those individuals surviving at least to 7 days.**

A and C: Kaplan-Meier survival estimates and 95% confidence intervals overlaid with the corresponding parametric survival curves fit to a Gompertz distribution. “+” indicates a censored data point. B and D: log(Hazard) (equivalent to log[likelihood of death]) curves for corresponding data fit to a Gompertz distribution; bands represent 95% confidence intervals. An increase in the intercept on this graph represents an increase in baseline mortality (α); an increase in the slope represents an increase in the rate of ageing (β). The unlikely conclusion of negative senescence in *H. melpomene* as demonstrated in B as well as the poor fit of survival data for this species to any hazard function as shown in Fig. S3 provided justification for the subsetting of the dataset for this species to only individuals surviving at least to 7 days for further analysis in this chapter. There was not similar support for performing this exclusion on any other species in this dataset, but C-D are plotted to show that doing so results in qualitatively similar conclusions (comparison with Fig. 1).

#### *2.9 Supplementary Note 9: Proposed scenario for the evolution of longevity in* Heliconius

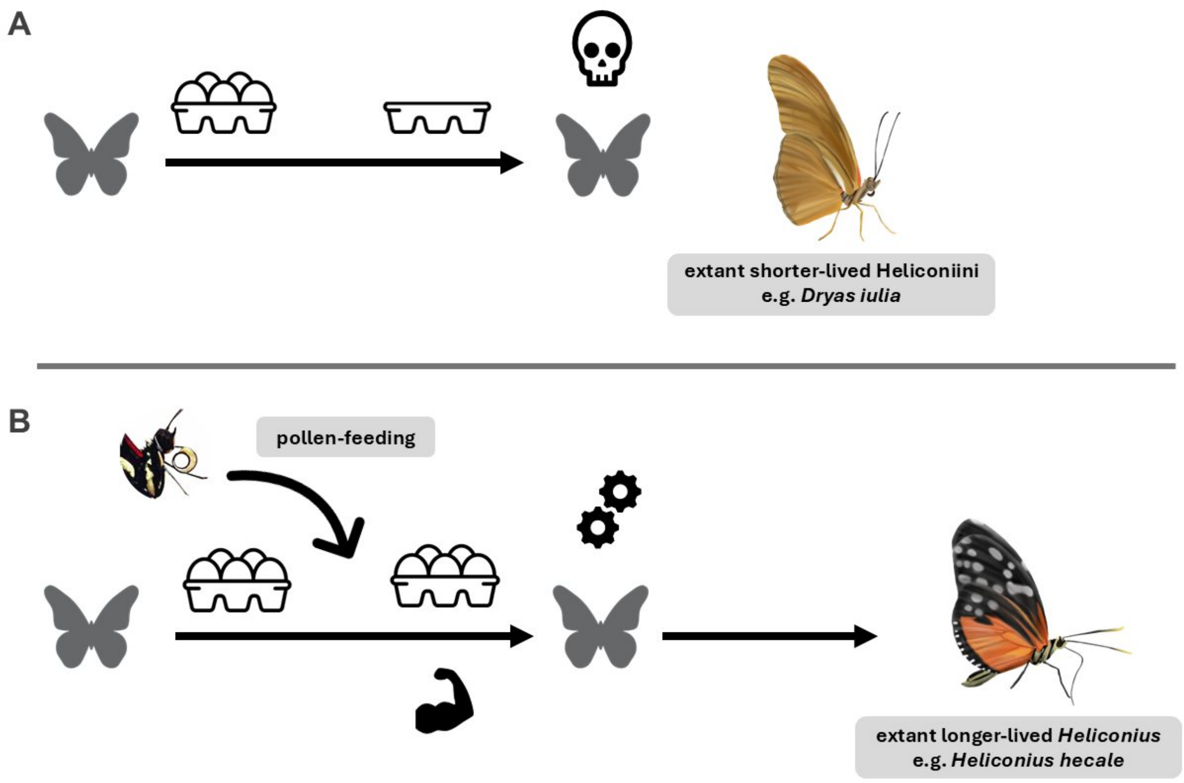

**Fig. S8: Proposed scenario for the evolution of longevity in *Heliconius*.**

**A** The last common ancestor of *Heliconius* and the outgroup Heliconiini was likely a non-pollen-feeder, and would have eclosed with a finite number of oocytes, reached the end of its reproductive lifespan after a few weeks, and died shortly thereafter. Longevity would not have been selected for in this instance because of this cap on reproductive output; this is the case we see in the extant shorter-lived Heliconiini, such as *Dryas iulia*. **B** The introduction of a pollen-feeding adaptation in the lineage leading to *Heliconius* would have conferred both direct fitness benefits (such as the reduced baseline mortality and enhanced grip strength shown in Figs. 2 and 3) as well as indirect fitness benefits, in the facilitation of a prolonged reproductive lifespan due to the continual production of oocytes. This would have allowed for selection on pro-longevity mechanisms and allowed for the evolution of longer life; this is the case we see in most extant *Heliconius*, such as *Heliconius hecale*. Heliconiini drawings are original artwork by Amaia Alcalde Antón and the *Heliconius* photograph is by Louise Bestea; all are reproduced here with permission.

#### References

1. Brewster, A.L.E. and G.W. Otis, *A Protocol for Evaluating Cost-Effectiveness of Butterflies in Live Exhibits.* Journal of Economic Entomology, 2009. **102**(1): p. 105-114.
2. Young, F.J., et al., *Enhanced long-term memory and increased mushroom body plasticity in* Heliconius *butterflies.* iScience, 2024: p. 108949.
3. Brown, K., *The biology of* Heliconius *and related genera.* Annual review of entomology, 1981. **26**(1): p. 427-457.
4. Lane, S.J., et al., *The effects of age and lifetime flight behavior on flight capacity in* Drosophila melanogaster*.* Journal of Experimental Biology, 2014. **217**(9): p. 1437-1443.
5. Sohal, R.S., *18 - Aging in Insects*, in *Biochemistry*, G.A. Kerkut and L.I. Gilbert, Editors. 1985, Pergamon: Amsterdam. p. 595-631.
6. Åhman, M. and B. Karlsson, *Flight endurance in relation to adult age in the green-veined white butterfly* Pieris napi*.* Ecological Entomology, 2009. **34**(6): p. 783-787.
7. Boggs, C.L., *REPRODUCTIVE ALLOCATION FROM RESERVES AND INCOME IN BUTTERFLY SPECIES WITH DIFFERING ADULT DIETS.* Ecology, 1997. **78**(1): p. 181-191.
8. Pásztor, K., et al., *Phenotypic senescence in a natural insect population.* Ecology and Evolution, 2022. **12**(12).
9. Davis, A.K., F.M. Smith, and A.M. Ballew, *A poor substitute for the real thing: captive-reared monarch butterflies are weaker, paler and have less elongated wings than wild migrants.* Biology Letters, 2020. **16**(4): p. 20190922.
10. R Core Team, *R: A Language and Environment for Statistical Computing*. 2023, R Foundation for Statistical Computing: Vienna, Austria.
11. Hebberecht, L., et al., *The evolution of adult pollen feeding did not alter postembryonic growth in* Heliconius *butterflies.* Ecology and Evolution, 2022. **12**(6).
12. Gompertz, B., *On the Nature of the Function Expressive of the Law of Human Mortality, and on a New Mode of Determining the Value of Life Contingencies.* Philosophical Transactions of the Royal Society of London, 1825. **115**: p. 513-583.
13. Williams, G.C., *Pleiotropy, Natural Selection, and the Evolution of Senescence.* Evolution, 1957. **11**(4): p. 398.
14. Therneau, T.M., *A Package for Survival Analysis in R.* 2023.
15. Therneau, T.M., *coxme: Mixed Effects Cox Models.* 2022.
16. Jackson, C., *flexsurv: A Platform for Parametric Survival Modeling in R.* Journal of Statistical Software, 2016. **70**(8): p. 1 - 33.
17. Hudson, D.W., et al., *Analysis of Lifetime Mortality Trajectories in Wildlife Disease Research: BaSTA and Beyond.* Diversity, 2019. **11**(10): p. 182.
18. Hess, K. and R. Gentleman, *muhaz: Hazard Function Estimation in Survival Analysis.* 2021.
19. Bronikowski, A. and T. Flatt, *Aging and its demographic measurement.* Nat Educ Knowl, 2010. **1**.
20. Wilson, D.L., *The analysis of survival (mortality) data: Fitting Gompertz, Weibull, and logistic functions.* Mechanisms of Ageing and Development, 1994. **74**(1): p. 15-33.
21. Colchero, F., O.R. Jones, and M. Rebke, *BaSTA: an R package for Bayesian estimation of age-specific survival from incomplete mark-recapture/recovery data with covariates.* Methods in Ecology and Evolution, 2012. **3**(3): p. 466-470.
22. Colchero, F. and J.S. Clark, *Bayesian inference on age-specific survival for censored and truncated data.* Journal of Animal Ecology, 2012. **81**(1): p. 139-149.
23. Revell, L.J., *phytools 2.0: an updated R ecosystem for phylogenetic comparative methods (and other things).* PeerJ, 2024. **12**: p. e16505.
24. Cicconardi, F., et al., *Evolutionary dynamics of genome size and content during the adaptive radiation of Heliconiini butterflies.* Nature Communications, 2023. **14**(1).

25. Bates, D., et al., *Fitting Linear Mixed-Effects Models Using lme4.* Journal of Statistical Software, 2015. **67**(1): p. 1-48.
26. Kuznetsova, A., P.B. Brockhoff, and R.H.B. Christensen, *lmerTest Package: Tests in Linear Mixed Effects Models.* Journal of Statistical Software, 2017. **82**(13): p. 1-26.
27. Kubinec, R., *Ordered Beta Regression: A Parsimonious, Well-Fitting Model for Continuous Data with Lower and Upper Bounds.* Political Analysis, 2022: p. 1-18.
28. Douma, J.C. and J.T. Weedon, *Analysing continuous proportions in ecology and evolution: A practical introduction to beta and Dirichlet regression.* Methods in Ecology and Evolution, 2019. **10**(9): p. 1412-1430.
29. Brooks, M.E., et al., *glmmTMB Balances Speed and Flexibility Among Packages for Zero-inflated Generalized Linear Mixed Modeling.* The R Journal, 2017. **9**(2): p. 378-400.
30. Hartig, F., *DHARMa: Residual Diagnostics for Hierarchical (Multi-Level / Mixed) Regression Models.* 2022.
31. Fox, J. and S. Weisburg, *An R companion to applied regression*. 3rd edition ed. 2019, Thousand Oaks, California: SAGE Publications, Inc. xxx, 577 pages.
32. Lüdecke, D., *sjPlot: Data Visualization for Statistics in Social Science.* 2023.
33. Dalbosco Dell'Aglio, D., O.W. Mcmillan, and S. Montgomery, *Using motion-detection cameras to monitor foraging behaviour of individual butterflies.* Ecology and Evolution, 2024. **14**(7).
34. Niitepõld, K. and C.L. Boggs, *Carry-over effects of larval food stress on adult energetics and life history in a nectar-feeding butterfly.* Ecological Entomology, 2022. **47**(3): p. 391-399.
35. Niitepõld, K., A. Perez, and C.L. Boggs, *Aging, life span, and energetics under adult dietary restriction in lepidoptera.* Physiol Biochem Zool, 2014. **87**(5): p. 684-94.
36. Pletcher, *Model fitting and hypothesis testing for age-specific mortality data.* Journal of Evolutionary Biology, 1999. **12**(3): p. 430-439.
37. Carroll, J., E. Korshikov, and T.N. Sherratt, *Post-reproductive senescence in moths as a consequence of kin selection: Blest's theory revisited.* Biological Journal of the Linnean Society, 2011. **104**(3): p. 633-641.
38. Carroll, J. and T.N. Sherratt, *Actuarial senescence in laboratory and field populations of Lepidoptera.* Ecological Entomology, 2017. **42**(5): p. 675-679.
39. Sielezniew, M., A. Kostro-Ambroziak, and Á. Kőrösi, *Sexual differences in age-dependent survival and life span of adults in a natural butterfly population.* Scientific Reports, 2020. **10**(1).
40. Watts, J.R. *Longevity studies in a tropical conservatory: are you getting your money's worth?* in *Invertebrates in Captivity Conference*. 2004. Sonoran Arthropod Studies Institute, Tucson, AZ.
